# Assessing the Potential of Switchgrass (*Panicum virgatum* L.) for Storing Carbon Belowground: Insights from a Multi-Site US Study

**DOI:** 10.1101/2025.08.01.668182

**Authors:** Eric W. Slessarev, Jennifer Pett-Ridge, Kyungjin Min, Asmeret A. Berhe, Srabani Das, Randall D. Jackson, Julie D. Jastrow, Megan Kan, Sandeep Kumar, Todd Longbottom, Karis J. McFarlane, Erik Oerter, Brian K. Richards, G. Philip Robertson, Gregg R. Sanford, Erin E. Nuccio

## Abstract

Agricultural intensification depletes belowground carbon (C) stocks, partly due to the exclusion of deep-rooted perennial species from landscapes where they were once dominant. Reintroducing deep-rooted perennials to cultivated land may help to mitigate SOC loss and restore ecosystem function. We quantified the effect of replacing shallow-rooted annual crops with a deep-rooted perennial grass, switchgrass (*Panicum virgatum* L.), by comparing 10 to 30 year-old switchgrass stands with paired annual row crop fields across the central and eastern USA. We hypothesized that switchgrass would store more root C and SOC than neighboring shallow-rooted annual crops, and that these effects would extend deeper than 30 cm. We also evaluated whether switchgrass stimulates decomposition of SOC at depth using radiocarbon (^14^C) to quantify replacement of slow cycling isotopically depleted SOC. Finally, we explored whether the effect of switchgrass on SOC is moderated by soil chemical and physical properties. While the effect of switchgrass on SOC in the surface 100 cm was positive at most sites, the average effect was highly uncertain and statistically indistinguishable from zero (mean difference in SOC = 0.6 kg C m^-2^ [95% CI −0.8 to +1.9 kg C m^-2^]). By contrast, we found that root biomass C was consistently more deeply distributed and more abundant under switchgrass, yielding an estimated additional 0.6 kg C m^-2^ in the surface 100 cm of soil under switchgrass [95% CI +0.5 to +0.7 kg C m^-2^]. ^14^C measurements suggested that root C inputs were adding to existing SOC without stimulating decomposition. The effect of switchgrass on belowground C was not strongly related to any of the environmental factors that we evaluated. Our observations show that root biomass C can contribute substantially to belowground C stocks when deep-rooted perennial grasses replace shallow-rooted annual crops.

## 1. Introduction

Agricultural expansion has caused substantial loss of belowground C in terrestrial ecosystems, contributing to the rise of atmospheric CO_2_ (Ogle et al., 2005; Sanderman et al., 2017; Padarian et al., 2022; Beillouin et al., 2023). Mitigating this belowground C loss is a major challenge facing modern agriculture (Bossio et al., 2020). One aspect of this challenge is finding ways to increase root C inputs. Roots transfer photosynthate belowground, where it is stored in root biomass, transferred to symbionts, or released into the soil as exudates, becoming soil organic carbon (SOC). Root C is transformed into comparatively slow-cycling mineral-associated SOC more efficiently than C inputs from aboveground plant biomass (Rasse et al., 2005; Austin et al., 2017; Sokol et al., 2019). Given that roots are an important conduit for C, the replacement of deep-rooted plants with shallow-rooted crops likely contributes to SOC loss (Hauser et al., 2022). Enhancing root C inputs by planting deep-rooted native vegetation, food crops, or biomass crops might help mitigate or reverse SOC loss (Kell, 2012; Lynch and Wojciechowski, 2015). Here we investigate the potential for enhancing root inputs and SOC with a specific emphasis on deep-rooted perennial grasses.

Grasses develop fibrous roots that typically are most abundant in the uppermost 15 cm of soil but can extend much deeper, particularly under water-limited conditions (Weaver, 1919). Many factors moderate belowground C allocation by grasses in addition to water stress. Perennial warm-season grasses native to North American prairie tend to allocate more C belowground than introduced forage grasses and biomass crops, which have been selected for aboveground production under relatively high resource conditions (Wilsey and Polley, 2006; Sprunger et al., 2017). Disturbances such as mowing and fire can enhance grass root production (Kitchen et al., 2009), but grazing that is too intense or frequent can also reduce root production (Biondini et al., 1998). The life history strategy of grasses also moderates root C inputs. Perennial species can provide year-round vegetative cover, which reduces or eliminates C loss associated with agricultural soil disturbance and ensures a more continuous supply of root C (King and Blesh, 2018). Native grasses that are both deep-rooted and perennial might therefore be particularly well-suited for restoring SOC.

Several comparative studies support the idea that deep-rooted perennial grasses enhance SOC. A global synthesis found that grass restoration enhances SOC (Conant et al., 2017). In Germany, a broad survey found that perennial grasslands store more SOC than neighboring land used to grow annual crops (Poeplau et al., 2021). Across the upper Midwest US, fields planted with switchgrass (*Panicum virgatum* L.) contained on average 7.74 Mg C ha^-1^ more SOC in the surface 60 cm of soil than fields planted with shallow-rooted annuals (Liebig et al., 2005). At two restored prairie sites in the north-central US, gains in SOC were observed at the surface, but losses occurred at depth (Matamala et al., 2008; Dietz et al., 2024). These studies show that perennial grasses with deep roots generally—but not always—enhance SOC below 30 cm.

Most studies of SOC change following establishment of perennial grasses do not specifically quantify increases in root biomass at depth or relate root investment to SOC dynamics, although there are exceptions (Matamala et al., 2008; Von Haden and Dornbush, 2017; Peixoto et al., 2022; Moore et al., 2025). Furthermore, root biomass often is not distinguished from other forms of coarse particulate SOC (Hirte et al., 2017). It thus remains unclear how much, and to what depth, planting deep-rooted perennials influences rooting depth distributions, or how roots relate to changes in deep SOC. Quantifying the influence of deeply rooted perennials at depth is important given that land management effects can extend well below the 30 cm depth often used as a reference point when studying SOC dynamics (VandenBygaart et al., 2011; Balesdent et al., 2018; Sulman et al., 2020; Skadell et al., 2023). Furthermore, there is increasing evidence for directional change in deep SOC due to warming and other global change factors (Ofiti et al., 2021; Soong et al., 2021; Hicks Pries et al., 2023), which suggests that understanding deep SOC dynamics is important for predicting the response of ecosystems to future conditions.

Deep roots may stimulate decomposition of existing SOC (Fontaine et al., 2007; Kuzyakov, 2010), so that that the effect of increased root biomass on SOC might not be strictly proportionate or positive. Root C inputs can increase microbial activity, accelerating decomposition of SOC (Helal and Sauerbeck, 1986; Huo et al., 2017). Stimulation of decomposition by roots is known as rhizosphere priming, and typically is quantified in soil microcosms and greenhouse experiments using isotope tracers (Huo et al., 2017). In the field, both ^13^C and ^14^C isotopes can be used to identify accrual of fresh root-derived C at depth (Slessarev et al., 2020). C isotopes are thus useful for evaluating the time-integrated effects of roots on deep SOC stocks. More specifically, shifts in SOC ^14^C content can be used to infer accelerated SOC turnover under increased C input rates (Stoner et al., 2021). The extent to which deep roots accelerate SOC turnover under field conditions is poorly constrained.

Another major unknown is how environmental factors moderate the effect of cropland perennialization on SOC storage. In a global synthesis study including 709 observations of SOC change following perennial cultivation (including grasses and tree crops), Ledo et al., (2020) observed that environmental factors like temperature and soil clay content explained only 20% of the variability in SOC dynamics, suggesting that a more complex set of controls were at play. In particular, soil geochemical properties influence the strength of organo-mineral association, and hence may moderate SOC accrual (Slessarev et al., 2022a). For instance, in grassland ecosystems, SOC stocks are positively correlated with exchangeable Ca^2+^ (O’Brien et al., 2015; Rasmussen et al., 2018; Von Fromm et al., 2021; King et al., 2023). Exchangeable cations help link organic matter to mineral surfaces (Rowley et al., 2018; Shabtai et al., 2023). For instance, at one site in southern Michigan, exchangeable Ca^2+^ moderated SOC accrual following afforestation (Morris et al., 2007). The broader role of exchangeable cations and related soil chemical properties in moderating SOC dynamics following the establishment of perennial grasses has not been explored.

We investigated the effects of replacing shallow-rooted annual crops with switchgrass, a deep-rooted perennial grass native to the North American tallgrass prairies. We asked three main questions: (1) To what depth does planting switchgrass increase root biomass, and are increases in root biomass accompanied by changes in the quantity of SOC at depth? (2) Are shifts in C isotopic composition after planting switchgrass more dramatic than shifts in total SOC, suggesting enhanced SOC turnover? (3) Do soil properties, including exchangeable Ca^2+^, moderate the effect of switchgrass on SOC?

To address these questions, we made paired comparisons of croplands planted with shallow-rooted crops (e.g., corn, wheat, soybeans) and 10 to 30 year-old switchgrass stands at 12 sites across the central and eastern USA. We hypothesized that switchgrass would yield higher root biomass and greater SOC than neighboring shallow-rooted crops, with these effects extending below 30 cm. We used C isotopes to evaluate the possibility that roots enhanced SOC turnover at depth. Finally, we hypothesized that differences in SOC under the two vegetation types would be most dramatic at sites with finer textured soils with high concentrations of exchangeable Ca^2+^. By testing these hypotheses, this work advances our understanding of how and where perennial grasses can restore SOC in cropland soils.

## 2 Methods

### 2.1 Study design and field sites

Our study was based on a paired-plot sampling design, in which mature plots planted with switchgrass were compared to neighboring plots planted with shallow-rooted annual crops. The advantage of this sampling design is tractability: paired-plot surveys can be used to evaluate land management effects rapidly across a wide range of environmental conditions (Liebig et al., 2005; Assad et al., 2013; Hong et al., 2020; Mosier et al., 2021; Becker et al., 2022). Differences in SOC between paired treated and control plots can be thought of as “relative changes”. Relative changes reflect the difference between treated and control systems at a single moment in time but yield no direct information about actual SOC dynamics (Sanderman and Baldock, 2010; Olson et al., 2014). Furthermore, the paired plot sampling design assumes that treatment and control plots initially had identical soil properties. Deviations from this assumption add to uncertainty.

We selected twelve sites that spanned a wide range of temperatures, precipitation regimes, and geologic settings and supported different switchgrass cultivars (Table 1). At each site we collected three replicate 2.5 m soil cores under each vegetation type, or until the depth of refusal. We attempted to sample the two vegetation types at closely paired or interspersed locations. When this was not possible, soil in adjacent (100 to 1000 m distant) fields with identically mapped soil series were sampled in triplicate. At two sites featuring long-term agricultural experiments, we sampled switchgrass and neighboring corn plots within interspersed treatment blocks. These sites included site 9, the Great Lakes Bioenergy Research Center’s Biofuel Cropping System Experiment in Hickory Corner, MI (Sprunger and Robertson, 2018) and site 11, the Wisconsin Integrated Cropping Systems Trial in Arlington, WI (Sanford et al., 2012). Sites 8, 10, and 12 at Cornell University, Ithaca, NY (Das et al., 2018), Fermilab, Batavia, IL (Adkins et al., 2019; Kelly-Slatten et al., 2023), and Bristol, SD (Lai et al., 2018; Min et al., 2021) featured replicate switchgrass plots but no replicate shallow-rooted annual crop plots; in these cases, we sampled adjacent shallow-rooted annual crop fields. At all other sites (sites 1-7), switchgrass and shallow-rooted annual crop plots were single fields that we subsampled. At sites 4 and 6 in Texas and North Carolina, we were nonetheless able to sample paired locations across field boundaries.

**Table 1.**
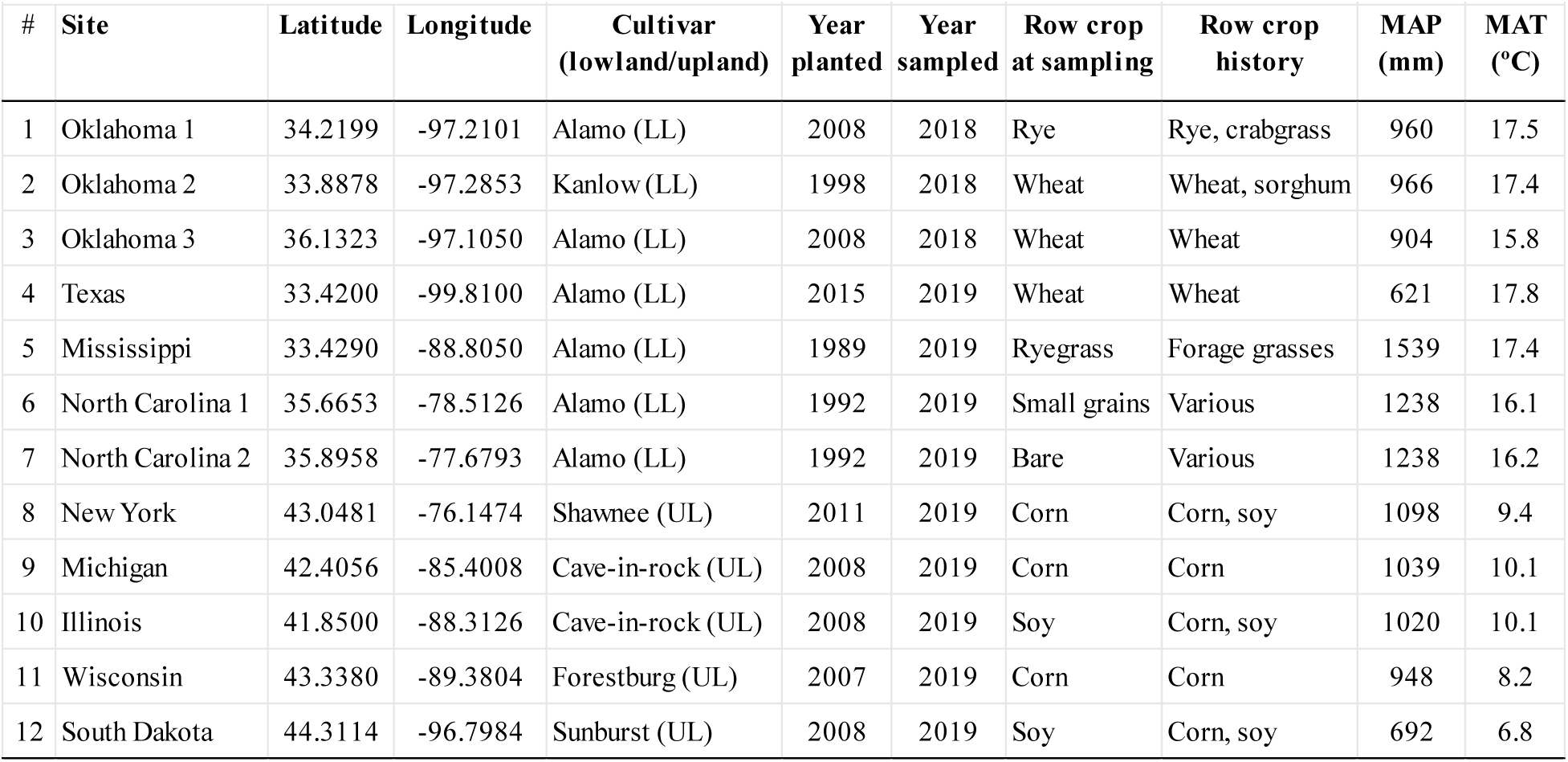
Description of 12 field sites featuring paired switchgrass and shallow-rooted annual crops. . Sites 1-3 were managed by the Noble Research Institute; Site 4 was managed by the Bud Smith Plant Material Center (USDA); site 5 by Mississippi State University; sites 6-7 by the North Carolina State Piedmont and Coastal Plains Agricultural Research Stations; site 8 by Cornell University; site 9 by the Great Lakes Bioenergy Research Center; site 10 by Argonne National Laboratory together with Fermi National Accelerator Laboratory; site 11 by the Wisconsin Integrated Cropping Systems Trial team at the University of Wisconsin-Madison; and site 12 by South Dakota State University. The column “shallow-rooted annual crop at sampling” designates the plant cover during the sampling visit, whereas “shallow-rooted annual crop history” summarizes shallow-rooted annual crop field cover over the long term. Climate data are averages for 2009-2018 derived from gridMET (Abatzoglou, 2013).

| Table 1. Description of field sites |  |  |  |  |  |  |  |  |  |  |
| --- | --- | --- | --- | --- | --- | --- | --- | --- | --- | --- |
| # | Site | Latitude | Longitude | Cultivar<br>(lowland/upland) | Year<br>planted | Year<br>sampled | Row crop<br>at sampling | Row crop<br>history | MAP<br>(mm) | MAT<br>(°C) |
| 1 | Oklahoma 1 | 34.2199 | -97.2101 | Alamo (LL) | 2008 | 2018 | Rye | Rye, crabgrass | 960 | 17.5 |
| 2 | Oklahoma 2 | 33.8878 | -97.2853 | Kanlow (LL) | 1998 | 2018 | Wheat | Wheat, sorghum | 966 | 17.4 |
| 3 | Oklahoma 3 | 36.1323 | -97.1050 | Alamo (LL) | 2008 | 2018 | Wheat | Wheat | 904 | 15.8 |
| 4 | Texas | 33.4200 | -99.8100 | Alamo (LL) | 2015 | 2019 | Wheat | Wheat | 621 | 17.8 |
| 5 | Mississippi | 33.4290 | -88.8050 | Alamo (LL) | 1989 | 2019 | Ryegrass | Forage grasses | 1539 | 17.4 |
| 6 | North Carolina 1 | 35.6653 | -78.5126 | Alamo (LL) | 1992 | 2019 | Small grains | Various | 1238 | 16.1 |
| 7 | North Carolina 2 | 35.8958 | -77.6793 | Alamo (LL) | 1992 | 2019 | Bare | Various | 1238 | 16.2 |
| 8 | New York | 43.0481 | -76.1474 | Shawnee (UL) | 2011 | 2019 | Corn | Corn, soy | 1098 | 9.4 |
| 9 | Michigan | 42.4056 | -85.4008 | Cave-in-rock (UL) | 2008 | 2019 | Corn | Corn | 1039 | 10.1 |
| 10 | Illinois | 41.8500 | -88.3126 | Cave-in-rock (UL) | 2008 | 2019 | Soy | Corn, soy | 1020 | 10.1 |
| 11 | Wisconsin | 43.3380 | -89.3804 | Forestburg (UL) | 2007 | 2019 | Corn | Corn | 948 | 8.2 |
| 12 | South Dakota | 44.3114 | -96.7984 | Sunburst (UL) | 2008 | 2019 | Soy | Corn, soy | 692 | 6.8 |

We selected switchgrass stands that were greater than five years old (Table 1). In most cases, the age of the switchgrass stand was equal to the time since land management diverged under switchgrass and adjacent shallow-rooted annual crop fields, which we term the “contrast age”. At site 9 in Illinois, switchgrass plots had been a long-term row crop field but were planted with cool-season grasses from 1971-2007 (*Bromus inermis*, *Poa* species, and volunteer *Agropyron repens*). We set the contrast age to 48 years at this site, since vegetation diverged in 1971. At site 4 in Texas, faulty records led us to sample a wheat field that had been previously planted with switchgrass and then converted back to wheat cultivation four years before our visit. In this case, we assigned a contrast age of four years, given that this was the amount of time that had elapsed since land management diverged.

Management variables were only loosely controlled across sites. In particular, the shallow-rooted annual crop type could not be matched across sites given the diversity of cropping systems across the study region; however, we sampled corn-soy or continuous corn cropping systems when these were available for comparison. At all but one site (site 6 in North Carolina), switchgrass plots were unfertilized. The fertilization rates and tillage practices applied to the shallow-rooted annual crops were not constrained at most sites; hence, we made no attempt to control for these variables in our analyses.

### 2.2 Field sampling protocol

We collected three deep soil cores under each vegetation type at each site (3 replicates * 2 vegetation types * 12 sites = 72 cores). Sites 1-3 in Oklahoma were part of a pilot study (Slessarev et al., 2020), and cores were collected using a Giddings probe (10.2 cm diameter). At sites 4-12 we collected cores using a Geoprobe 54LT direct push coring rig fitted with either MC5 tooling (3.81 cm diameter; sites 4-8) or MC7 tooling (7.62 cm diameter, sites 9-12). Cores were taken directly under switchgrass plants or shallow-rooted annual crop plants in order to maximize our ability to quantify rooting depth distributions; consequently, all C stock estimates reported here apply only to the area adjacent to plants and should not be used to extrapolate to the field scale. Furthermore, in some cases, shallow-rooted annual crop fields were fallow or bare when sampling occurred, in preparation for planting. The root biomass estimates were collected during the vegetative growth stage prior to flowering (June-July) and are hence only a seasonal snapshot and do not represent average biomass throughout the year.

Cores were sampled to a depth of at least 2 m, except for several cases when subsurface barriers or mechanical problems prevented coring beyond 100 cm. All cores were sectioned immediately in the field for root collection. Cores at sites 1-3 were sectioned at fixed depths (30 cm). Cores at sites 4-12 were sectioned in 20 cm increments for the first three depth intervals and in 30 cm increments thereafter, except when soils had strongly expressed horizons, in which case the breaks between sampling increments were shifted to accommodate horizon boundaries. Soil samples were weighed in the field, roots were removed, and soils were homogenized by hand and then subsampled for gravimetric water content analysis. Water content subsamples were immediately stored in Mylar® bags and heat sealed; these were shipped to Lawrence Livermore National Laboratory for a separate water-isotope study (Oerter et al. 2021).

Within 48 h of sampling in the field, we manually collected live roots in every soil core increment by removing them from the soil with forceps. In the uppermost two sampling increments, we limited root collection to 10 person-min per sample. Roots were later washed in deionized water in the lab, stored in paper envelopes, and dried at 60 °C for weighing and elemental analysis. The root biomass collected from the surface sample at site 4, core 1, was lost in a windstorm; in this case we imputed a value by averaging the other two replicate cores.

### 2.3 Soil analyses

Soil samples were stored in a cooler and then shipped overnight within 3 days to Lawrence Livermore National Laboratory for physical and chemical analyses. We then air dried, weighed, and sieved soil samples to 2 mm. Prior to soil C and isotope analysis, visible root pieces (including those that passed through the 2 mm sieve) were removed with forceps and combined with the roots we collected in the field. Coarse material (> 2 mm) was weighed, and its volume was estimated by water displacement in a graduated cylinder. All analyses were carried out on the < 2 mm fraction. We submitted subsamples of this fraction to the Oregon State University soil analytical laboratory for physical and chemical analyses. These included soil particle size (pipette method), soil pH in a 1:1 soil:water mixture, and exchangeable Ca, Mg, K, Na, and Al via 1 M BaCl_2_ extraction. Soil pH was measured on all but two depth intervals (n = 627), whereas texture and exchangeable cation measurements were generally limited to the upper 150 cm of each core (n = 430).

Samples with a pH > 6.5 were screened for inorganic C by mixing 1 M phosphoric acid with a finely ground sample in a sealed glass jar fitted with a septum and measuring the change in headspace pCO_2_. If they contained detectable inorganic C, these samples were then treated with 1 M HCl until inorganic C was removed (Slessarev et al., 2020). While this approach may result in small amounts of SOC oxidation, it is the only reliable way to prepare samples for C isotope analysis on the organic fraction, particularly for ^14^C (Harris et al., 2001).

Samples were prepped for C isotope analysis by pulverizing 10 g subsamples in a roller mill. We then submitted this finely-ground material to the UC Berkeley Stable Isotope Facility for ^13^C analysis (https://nature.berkeley.edu/stableisotopelab/). The ^13^C content of each sample (δ^13^C) was reported on a per mil (‰) basis relative to the V-PDB isotope standard. We used total C values reported from this analysis when calculating organic carbon concentrations and stocks because these samples were acid treated and hence assumed to be carbonate free; this provided a consistent basis for ^13^C, ^14^C, and total C analyses. All samples were analyzed for total organic C and ^13^C (n = 629).

Radiocarbon values were measured on the NEC 1.0 MV Tandem accelerator mass spectrometer (AMS) or the FN Tandem Van de Graaff AMS at the Center for AMS at Lawrence Livermore National Laboratory. Samples were prepared for ^14^C measurement by sealed-tube combustion to CO_2_ in the presence of CuO and Ag and then reduced onto iron powder in the presence of H_2_ (Vogel, Southon, Nelson, & Brown, 1984). The ^14^C content of each sample (Δ^14^C) was reported in ‰ relative to the absolute atmospheric ^14^C activity in 1950. To calculate Δ^14^C, measured δ^13^C values were used to correct for mass-dependent fractionation to yield ^14^C activity at a reference δ^13^C of −25‰ (Stuiver & Polach, 1977). Because sample collection and analysis occurred within a short period, no correction was performed for decay of ^14^C between sampling and analysis. The average instrument uncertainty for Δ^14^C was ±2.6‰. Radiocarbon analyses were generally limited to the upper 150 cm of each soil core (n = 456).

### 2.4 Equivalent mass calculations

We summarized the data from each soil core on a mineral mass equivalent basis (Gifford and Roderick, 2003; Wendt and Hauser, 2013; Rovira et al., 2015; Von Haden et al., 2020). The mass of mineral soil < 2 mm in each sampled interval was obtained by correcting the mass of each sample measured in the field for its moisture content and subtracting out the mass of particles > 2 mm and the mass of organic matter [mass OM = 2*mass OC (Pribyl, 2010)]. To calculate C stocks at pre-defined soil mass equivalents we fit a monotone cubic spline to predict cumulative C mass as a function of cumulative < 2 mm mineral mass on a core-by-core basis.

We calculated C stocks at equivalent masses of 420 kg m^-2^, 840 kg m^-2^, and 1400 kg m^-2^, which correspond to approximately 30-cm, 60-cm, and 100-cm depths on average across the dataset. Additional soil variables related to pH, exchangeable cations, and texture were summarized by calculating mass-weighted average values for each mass interval. When calculating mass-weighted averages, we weighted the value for each layer by its relative contribution to the mass interval being considered. For layers that were bisected by the upper or lower boundary of each mass interval, we only counted the contribution from the mass that fell within the interval.

Similarly, C isotope values (Δ^14^C, δ^13^C) were summarized at the specified mass intervals by computing C stock weighted averages. In these cases, values for each layer were weighted by the contribution of the layer’s C stock to the total C stock of the mass interval being considered.

Where layers were bisected by the upper or lower bounds of the mass interval, only the component of the C stock included in the interval was used for weighting, with the partial contribution determined using the spline function for cumulative carbon mass.

### 2.5 Root quantification

We quantified the total root mass in each depth interval in mg root per g of < 2 mm sieved soil. As described in Section 2.2, roots were manually removed in the field using forceps, washed in the laboratory, and dried at 60 °C. Root masses were then used to compute total root inventories at the cumulative soil masses specified above. Additionally, for each soil core, we computed the total root mass obtained to the maximum sampling depth and normalized root inventories by this value to obtain the cumulative fraction of total roots sampled as a function of depth. We then used linear interpolation to find the depth at which 95% of roots were encountered. This value is analogous to the “D_95_” parameter used to summarize maximum rooting depths, but it is based on observed roots [i.e., we did not extrapolate rooting distributions as in (Schenk and Jackson, 2005)]. We also estimated root C stocks by measuring the total C content of the roots collected at each site at the UC Berkeley Stable Isotope Facility. Not all samples provided sufficient root material for total C analysis; therefore, when calculating root C stocks, we assumed the C content of unmeasured roots was approximated by the average value across all sites (37% C).

### 2.6 Isotope mixing model

We used ^14^C as a tracer of newly added C deeper in the soil column. Below the uppermost 10-20 cm of soil, SOC tends to be ^14^C depleted relative to the atmosphere. Hence, if recently fixed atmospheric C is added to deep soil, it will register as an increase in the Δ ^14^C value (Slessarev et al., 2020). We estimated the fraction of SOC derived from this additional recent C at depth under switchgrass:

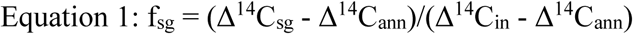

Where f_sg_ represents the fraction of additional recent SOC under switchgrass, Δ^14^C_sg_ is the Δ^14^C value of the soil under switchgrass, Δ^14^C_ann_ is the Δ^14^C of the soil under the shallow-rooted annual crop, and Δ^14^C_in_ is the Δ^14^C value of plant C inputs since planting of switchgrass.

Δ^14^C_in_ could not be observed directly. Instead, we obtained Δ^14^C_in_ by modeling the Δ^14^C of a hypothetical C pool governed by a first-order decay process (Slessarev et al. 2020). Δ^14^C_in_ depended only on specifying a decay rate constant for the hypothetical C pool. Because the true decay rate was unknown, we assumed two extreme cases: a fast decay rate [one year half-life, k = ln(2)] and slow decay [10-year half-life, k = 0.1*ln(2)]. We then assumed that Δ^14^C_in_ was equal to the average of these two extremes. Δ^14^C_in_ was modelled separately at each site based on the duration since switchgrass planting and the time-dependent concentration of ^14^C in the Northern Hemisphere atmosphere using the R package soilR (Sierra et al., 2012).

While we also used ^13^C as a qualitative tracer of switchgrass C, we refrained from applying a mixing model to ^13^C because the δ^13^C value of C sources could not be well constrained at most of our field sites. The δ^13^C of C inputs under the shallow-rooted annual crop was hard to constrain because both C_3_ and C_4_ crops were often grown in rotation, with uncertain contributions from each crop type.

### 2.7 Texture-corrected SOC stocks

In practice, our ability to pair soil conditions was not perfect, and soil properties differed even in closely adjacent fields (e.g., within 100 m). We addressed these discrepancies by calculating texture-corrected SOC values at a site level. The corrected values were computed by assuming a site and soil mass increment specific linear relationship between SOC stocks and soil silt + clay stocks. We then corrected SOC levels to reflect the mean silt + clay stock at that site and soil mass increment:

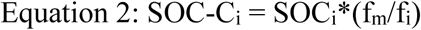

In this equation, SOC-C_i_ is the corrected SOC stock for core i, SOC_i_ is the uncorrected stock, f_i_ is the fine fraction (silt + clay) stock, and f_m_ is the mean silt + clay stock for the site and soil mass interval being analyzed. The correction factor accounts for initial differences in SOC stock under the following assumptions: (1) the majority of SOC was associated with silt and clay sized particles; (2) initial SOC loading values [SOC/(silt + clay)] were homogenous within a given site and mass increment such that all variability in initial SOC stocks was due to variability in silt + clay content (3) silt + clay content was static over the time since switchgrass establishment and not sensitive to management.

### 2.8 Statistical analysis

We quantified the effect of switchgrass on SOC, root biomass, and C isotopes with depth by calculating the mean difference in each property (switchgrass – shallow-rooted annual crop) across sites. We used a non-parametric bootstrap resampling approach to evaluate uncertainty when computing mean differences (Davison and Hinkley, 1997). We used this approach because it allowed us to represent the nested structure of our dataset and propagate both within-site and across-site uncertainty.

The bootstrapping procedure involved (1) resampling the three soil cores taken under each plant type at each site with replacement. At sites where we achieved core-level pairing (sites 4, 6, 9, and 11), we resampled pairs rather than cores with replacement; (2) computing the mean difference or other statistic for each property at each site using the resampled data; (3) resampling sites with replacement and computing the overall mean of each statistic across sites. This procedure was repeated a total of 5,000 times to estimate the sampling distribution for each statistic. We then calculated bias-corrected and accelerated 95% confidence intervals from this distribution. Bias-corrected values were obtained from bootstrap distributions using the R package “coxed” (Kropko and Harden, 2020).

We also computed a correlation statistic, Spearman’s ρ, to test whether differences in SOC or root stocks under the two vegetation types were related to environmental variables. We used this statistic because it makes few assumptions about the functional form of the relationship between two correlated variables (e.g., relationships need not be linear). We estimated confidence intervals for ρ using the bootstrapping procedure described above.

We first computed ρ to evaluate relationships between SOC stocks, root C stocks, δ ^13^C, and Δ^14^C values calculated at 1400 kg m^-2^, or approximately 100 cm depth. We then examined correlations between C stock variables and environmental variables. The environmental variables included soil chemical and physical indices that we hypothesized might affect SOC accrual: soil pH, silt + clay content, effective cation exchange capacity (ECEC), total exchangeable calcium, and the site-averaged SOC stock. ECEC was calculated by summing the charges of the exchangeable ions (Burt, 2014). We also included two climate indices (mean annual temperature and mean annual precipitation) and both the switchgrass cultivar and the vegetation contrast age. We note that relating differences in SOC to standing SOC stocks across all sites could create a spurious correlation due to the effect of regression to the mean (Slessarev et al., 2022b). We avoided this statistical artifact by relating changes in SOC between switchgrass and shallow-rooted annual crops to the overall site level mean rather than the mean of a particular vegetation cover (Lark et al., 2006).

In addition to confidence intervals, we selectively computed *p*-values to test specific null hypotheses that the correlation statistic, ρ, was equal to zero. We obtained *p*-values empirically via permutation tests by randomly shuffling the variable of interest without replacement and calculating ρ 5,000 times. We then computed *p*-values by dividing one plus the number of cases that exceeded the statistic of interest by one plus the total number of permutations (Davison and Hinkley, 1997).

## 3 Results

### 3.1 Variation in soil properties and SOC pools

The 12 sites varied widely in their soil properties (Table 2). For instance, the average pH of the surface ∼30 cm of soil ranged from pH 5.6 at site 6 in North Carolina, which featured highly weathered soils, to pH 8.1 at site 12 in South Dakota, where the soils formed on calcareous glacial till. SOC stocks, root C stocks, and C isotope values also varied widely across the 72 soil cores that we collected. Data from individual soil cores are presented together in Figure 2 to convey the variability in soil C pools and isotopic signatures across the 12 sites.

**Figure 1.**
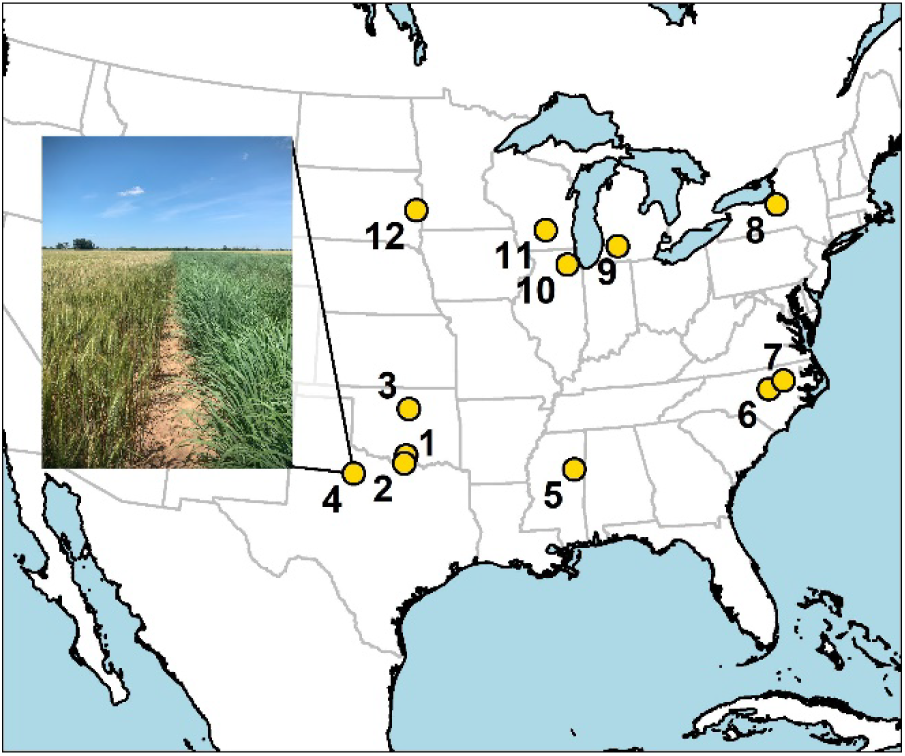
Map of study sites across the central and eastern USA. Numbering corresponds to site numbers listed in Table 1. Sites 1-7 were located in unglaciated landscapes and featured lowland switchgrass cultivars (e.g., Alamo). Sites 8-12 were located in glaciated landscapes and featured upland cultivars (e.g., Cave-in-Rock). Inset shows an example of spatial paring between switchgrass and shallow-rooted annual crop (wheat) at site 4 near Knox City, TX.

**Figure 2.**
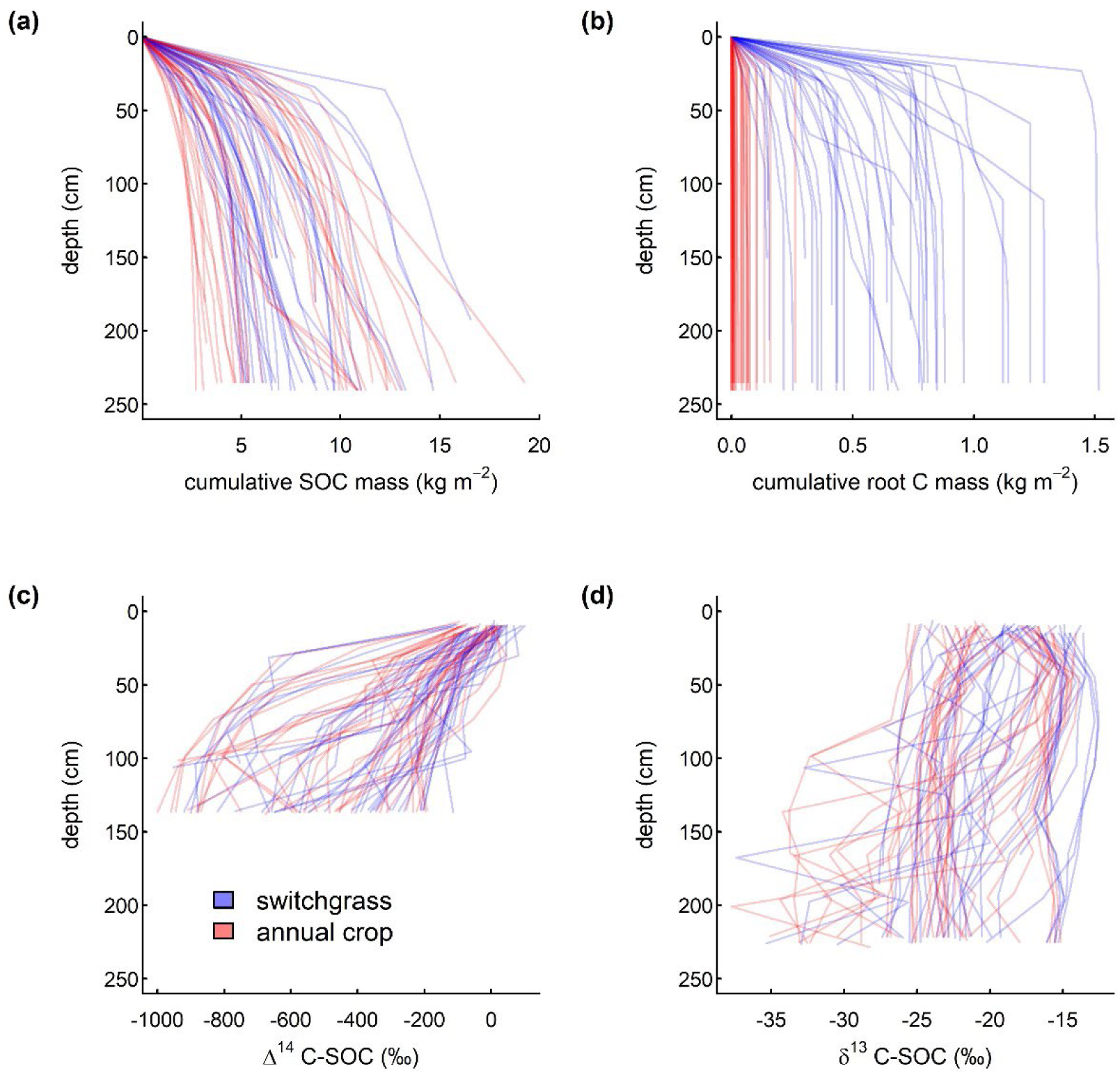
Variation in carbon pools and isotopes with depth at 12 sites in the central and eastern USA where annual crops were compared to adjacent mature switchgrass field. The four panels show (a) SOC, (b) root C, (c) ^14^C, and (d) ^13^C data pooled across all sites. Each line represents an individual soil core (n=72). At a few sites data were collected below 200 cm but are not shown here; ^14^C measurements terminated at 130 cm. Plots of these data disaggregated by site are included in Supplementary Information.

**Table 2.** Soil physico-chemical properties at twelve sites where shallow-rooted annual crops were compared to adjacent mature switchgrass fields. . Values represent mineral mass weighted averages of the upper 420 kg m^-2^ of soil, which is approximately equal to a depth interval of 0-30 cm. Data from switchgrass fields and shallow-rooted annual crop fields are averaged together. Values in parentheses are standard deviations. Ca-X, Mg-X, K-X, Na-X, and Al-X are exchangeable (BaCl_2_ extractable) Ca, Mg, Na, K and Al. ECEC is the effective cation exchange capacity.

| Table 2. Summary of soil physico-chemical properties |  |  |  |  |  |  |  |  |  |  |
| --- | --- | --- | --- | --- | --- | --- | --- | --- | --- | --- |
| Site # | pH | sand | silt | clay | Ca-X | Mg-X | K-X | Na-X | Al-X | ECEC |
|  |  | (%) |  |  | (cmol charge kg<sup>-1</sup>) |  |  |  |  |  |
| 1 | 6.2 (0.37) | 66 (6) | 26 (4) | 8 (2) | 3.6 (1.72) | 0.1 (0.06) | 1.6 (0.85) | 0 (-) | 0 (-) | 5.4 (2.55) |
| 2 | 7.0 (0.39) | 35 (9) | 41 (7) | 25 (2) | 9.9 (0.37) | 0.2 (0.05) | 3.2 (0.32) | 0 (-) | 0 (-) | 13.3 (0.51) |
| 3 | 6.5 (0.43) | 46 (10) | 17 (5) | 37 (12) | 7.7 (2.12) | 0.2 (0.03) | 6.7 (4.07) | 0.5 (0.88) | 0 (-) | 15 (6.52) |
| 4 | 7.3 (0.16) | 66 (2) | 17 (2) | 17 (2) | 5.8 (0.75) | 3.8 (0.44) | 0.3 (0.02) | 0.1 (0.03) | 0 (-) | 9.9 (1.14) |
| 5 | 5.8 (0.34) | 35 (13) | 39 (11) | 25 (4) | 10.5 (2.87) | 0.8 (0.21) | 0.2 (0.03) | 0.1 (0.06) | 0.5 (0.66) | 12.1 (2.52) |
| 6 | 5.6 (0.38) | 78 (15) | 8 (2) | 14 (16) | 1.1 (0.68) | 0.6 (0.27) | 0.2 (0.18) | 0 (0.01) | 0.1 (0.18) | 2 (0.98) |
| 7 | 5.7 (0.63) | 83 (6) | 12 (7) | 5 (2) | 0.5 (0.24) | 0.1 (0.14) | 0 (0.03) | 0 (0.01) | 0.4 (0.28) | 1 (0.32) |
| 8 | 6.4 (0.42) | 26 (3) | 44 (4) | 30 (4) | 10.3 (4.99) | 0.9 (0.41) | 0.1 (0.03) | 0.1 (0.16) | 0.1 (0.19) | 11.6 (5.26) |
| 9 | 6.1 (0.25) | 61 (20) | 20 (10) | 20 (11) | 6 (3.25) | 1.3 (0.63) | 0.2 (0.1) | 0.1 (0.02) | 0 (0.03) | 7.7 (3.76) |
| 10 | 5.7 (0.27) | 5 (3) | 62 (4) | 33 (6) | 12.1 (2.5) | 6 (1.5) | 0.3 (0.09) | 0.1 (0.01) | 0.1 (0.03) | 18.5 (3.75) |
| 11 | 6.3 (0.23) | 5 (1) | 64 (4) | 32 (4) | 9.3 (1.43) | 5.9 (1.27) | 0.3 (0.06) | 0.1 (0.01) | 0 (0.01) | 15.5 (2.7) |
| 12 | 8.1 (0.14) | 33 (13) | 31 (6) | 36 (10) | 10.5 (1.82) | 3.5 (1.21) | 0.2 (0.02) | 0.1 (0.02) | 0 (-) | 14.2 (2.79) |

SOC was most abundant near the surface, but SOC stocks continued to increase even as the depth sampled exceeded 100 cm, where stock values ranged from 2.5 kg C m^-2^ to 14 kg C m^-2^ (Fig. 2a). SOC stocks under the two vegetation types fell within the same broad range. By contrast, root C stocks were dramatically different under the two vegetation types (Fig. 2b).

Under shallow-rooted annual crops, the root C stock captured during sampling was generally confined to the uppermost 30 cm of soil and roots contributed < 0.1 kg C m^-2^, whereas under switchgrass, root C stocks ranged from ∼ 0.1 to > 1 kg C m^-2^.

Carbon isotopes were similar in SOC under the two vegetation types. SOC radiocarbon (^14^C) values were greatest and least variable near the surface and declined with depth, ranging from more than −200 ‰ to nearly −1000 ‰ at depths below 100 cm (Fig. 2c). These very low values may have been caused by the presence of petrogenic C (see Discussion). SOC ^13^C values varied in the surface 30 cm and were bimodally distributed, with one group of soil cores from sites 1-4, 10, and 11 yielding δ^13^C values between −17 and −13 ‰ and the remaining cores yielding values that ranged between −27 and −17 ‰. At depths greater than 30 cm, δ^13^C values were erratic and, in some cases, reached values less than −35 ‰ (Fig. 2d).

### 3.2 Effect of switchgrass on C stocks and isotopes

We compared SOC stocks and C isotope values under the two vegetation types at equivalent soil mineral masses of 0-420, 420-840, and 840-1400 kg m^-2^, which corresponded to depth intervals 0-30, 30-60, and 60-100 cm, on average. SOC stocks were higher under switchgrass in the 0-420 kg m^-2^ interval at 9 out of 12 sites (Fig. 3a). At the other depth intervals, SOC stocks were similar under the two vegetation types. Correcting for soil texture differences within each site (see Methods) changed this pattern; corrected SOC values were significantly greater under switchgrass at the surface (Fig. 3b). Mean SOC stocks were slightly higher under switchgrass in the uppermost 0-1400 kg of soil in both uncorrected and corrected cases, but the difference in SOC between switchgrass and shallow-rooted annual plots was not statistically distinguishable from zero. The uncorrected difference in SOC in the 0-1400 kg mass interval was 0.6 kg C m^-2^ [95% CI −0.8 to +1.9 kg C m^-2^], and the texture-corrected difference was 0.9 kg C m^-2^ [95% CI − 0.5 to +2.8 kg C m^-2^].

**Figure 3.**
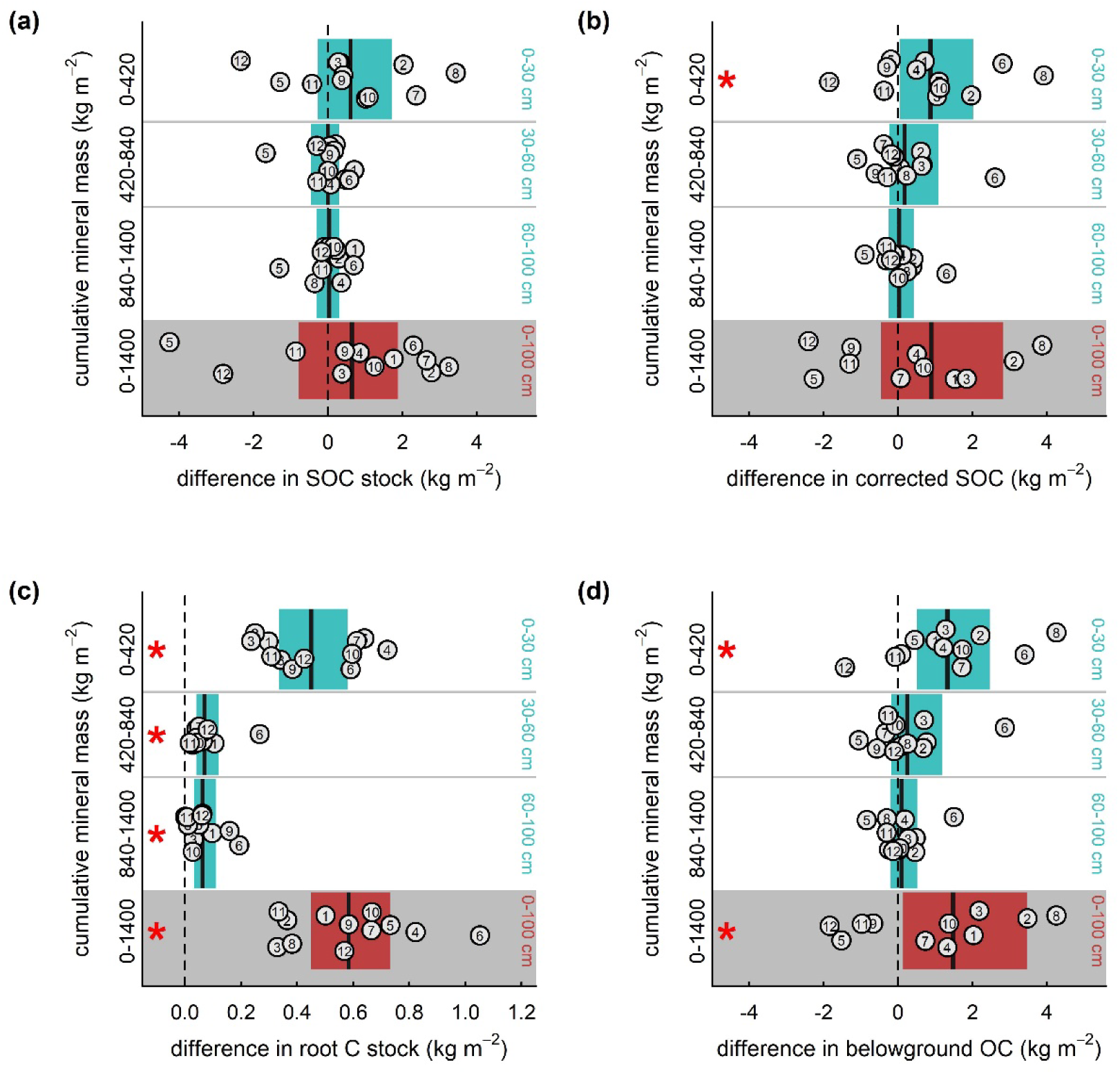
Differences in C stocks between shallow-rooted annual crop fields and adjacent mature switchgrass fields in the central and eastern USA, computed across soil mass intervals. Panels show differences (switchgrass – shallow-rooted annual crop) in SOC stocks (a), texture-corrected SOC stocks (b), root C stocks (c) and total belowground organic C stocks, including SOC and root C (d). Data are shown in horizontal bands at 0-420, 420-840, and 840-1400 kg m^-2^ mineral mass intervals, which correspond to approximately 0-30, 30-60, and 60-100 cm depth intervals. Data for the entire uppermost 0-1400 kg m^-2^ (approximately 0-100 cm) of soil are shown at the bottom of each panel. Points show observed mean differences at the twelve sites, with site numbers shown as in Table 1 and random noise added along the y axis to reduce overlap. Black vertical lines show the observed across-site mean at each interval, and shaded regions show bootstrapped 95% confidence intervals, with blue indicating interval-by-interval summaries and red indicating the whole profile to 1400 kg m^-2^. A red asterisk indicates depth intervals at which the confidence interval does not include zero.

In contrast to total SOC, root biomass C was consistently and significantly higher under switchgrass. The 95% confidence intervals for differences in root C were strictly positive at all soil intervals considered (Fig. 3c). Furthermore, the rooting depth—and not just the total amount of root C—was also greater under switchgrass. We estimated that the depth at which 95% of root biomass was observed was 92 cm [95% CI 70 to 116 cm] under switchgrass and 31 cm [95% CI 10 to 62 cm] under shallow-rooted annual crops. The mean difference in rooting depth was 62 cm [95% CI 31 to 88 cm].

We evaluated total belowground organic C storage by adding root C and texture-corrected SOC stocks. Total belowground organic C was higher under switchgrass in the 0-420 kg mass interval (1.3 kg C m^-2^ [95% CI 0.5 to 2.6 kg C m^-2^]). The effect of switchgrass on belowground C stocks at greater depths remained small and uncertain (Fig. 3d). Over the entire 0-1400 kg mass interval, the mean difference in belowground C was statistically significant and equaled 1.5 kg C m^-2^ [95% CI 0.1 to 3.5 kg C m^-2^].

Distinct C isotope signatures were detectable under switchgrass. Radiocarbon values under switchgrass and shallow-rooted annual crops were variable and broadly similar in the 0-420 and 420-840 kg depth intervals (i.e., the surface 0-60 cm), although differences tended to be positive at most sites, with the exception of one site (South Dakota) which featured substantially more ^14^C-depleted SOC under switchgrass at these depths (Fig. 4a). Deeper in the soil profile within the 840-1400 kg depth interval (approximately 60-100 cm), switchgrass Δ^14^C values were comparable to or higher than values under shallow-rooted annual crops at all but two sites. The difference in Δ^14^C within this depth range averaged +62 ‰ [95% CI +4 to +165 ‰]. When we calculated the overall shift in Δ^14^C across the uppermost 1400 kg of soil, we estimated that switchgrass increased Δ^14^C values by +43 ‰ [95% CI –7 to +105 ‰]. Unlike Δ^14^C, δ^13^C values were generally higher under switchgrass at all depths (Fig. 4b). Within the uppermost 1400 kg of soil, all but three sites had higher δ^13^C values under switchgrass, and the average shift in δ^13^C was 0.92 ‰ [95% CI 0.06 – 1.80 ‰].

**Figure 4.**
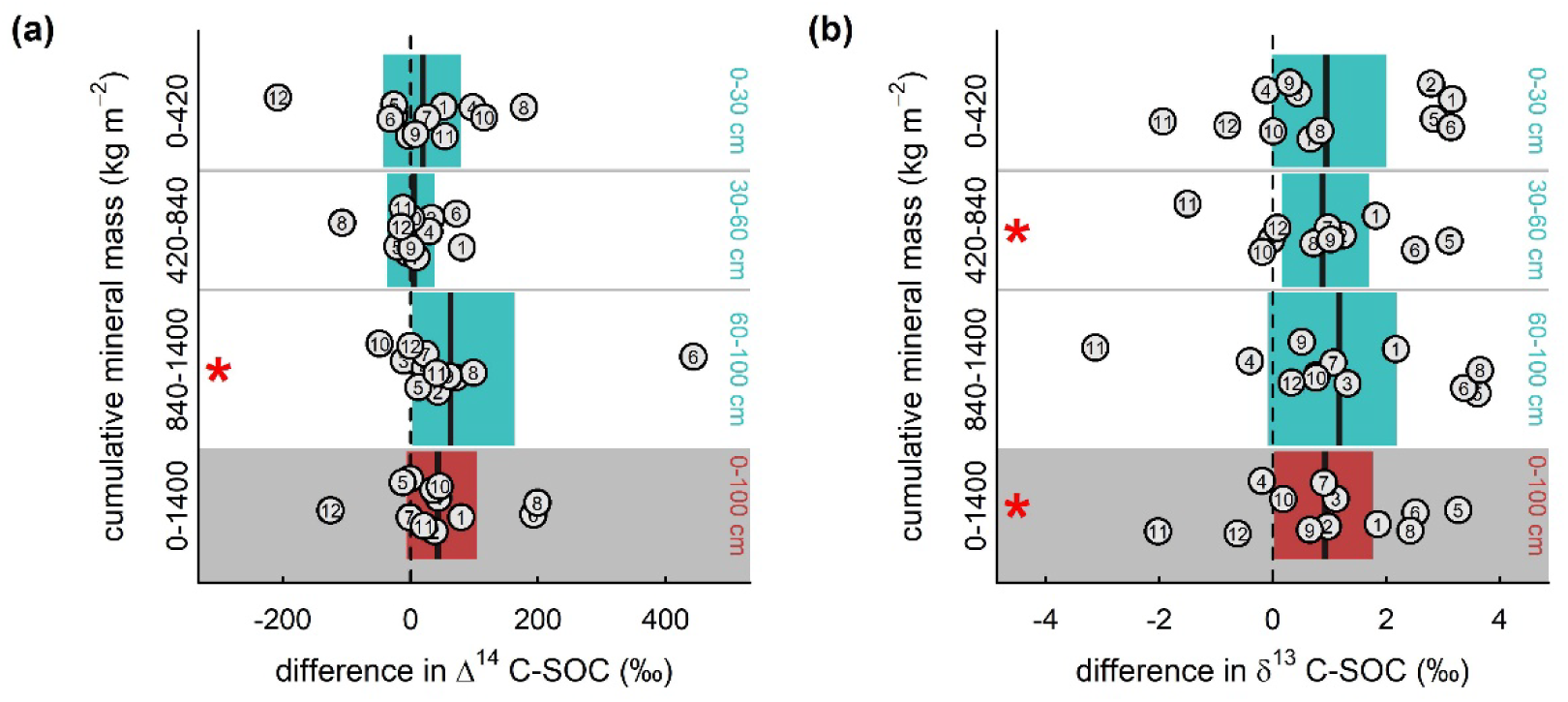
Differences in C isotopes between shallow-rooted annual crop fields and adjacent mature switchgrass fields in the central and eastern USA, computed across soil mass intervals. Panels show differences (switchgrass – shallow-rooted annual crop) in C-stock weighed average Δ^14^C (a) and δ^13^C (d). Data are shown in horizontal bands at 0-420, 420-840, and 840-1400 kg m^-2^ mineral mass intervals, which correspond to approximately 0-30, 30-60, and 60-100 cm depth intervals. Data for the entire uppermost 0-1400 kg m^-2^ (0-100 cm) of soil are shown at the bottom of each panel. Points show observed mean differences at the twelve sites with random noise added along the y axis to reduce overlap. Black vertical lines show the observed across-site mean at each interval, and shaded regions show bootstrapped 95% confidence intervals, with blue indicating interval-by-interval summaries and red indicating the whole profile to 1400 kg m^-2^. A red asterisk indicates depth intervals at which the confidence interval does not include zero.

### 3.3 Correlations between SOC dynamics and environmental variables

We explored relationships between environmental variables and the effect of switchgrass on SOC and root C across the twelve sites. The effect of switchgrass on texture-corrected SOC was not correlated with the effect of switchgrass on root C (ρ = −0.11 [95% CI –0.68 to +0.53]). The only strong correlation that we identified between the responses of C pools and isotopes to switchgrass was between the relative change in SOC and the relative change in Δ^14^C (ρ = 0.78, 95% CI +0.06 to +0.91) (Fig. 5a). Correlations between the effect of switchgrass on SOC and soil physico-chemical and environmental variables were weak in all cases, with ρ values between −0.5 and 0.5. Similarly, the effect of switchgrass on root C was not strongly correlated with any other variable (Fig. 5b).

**Figure 5.**
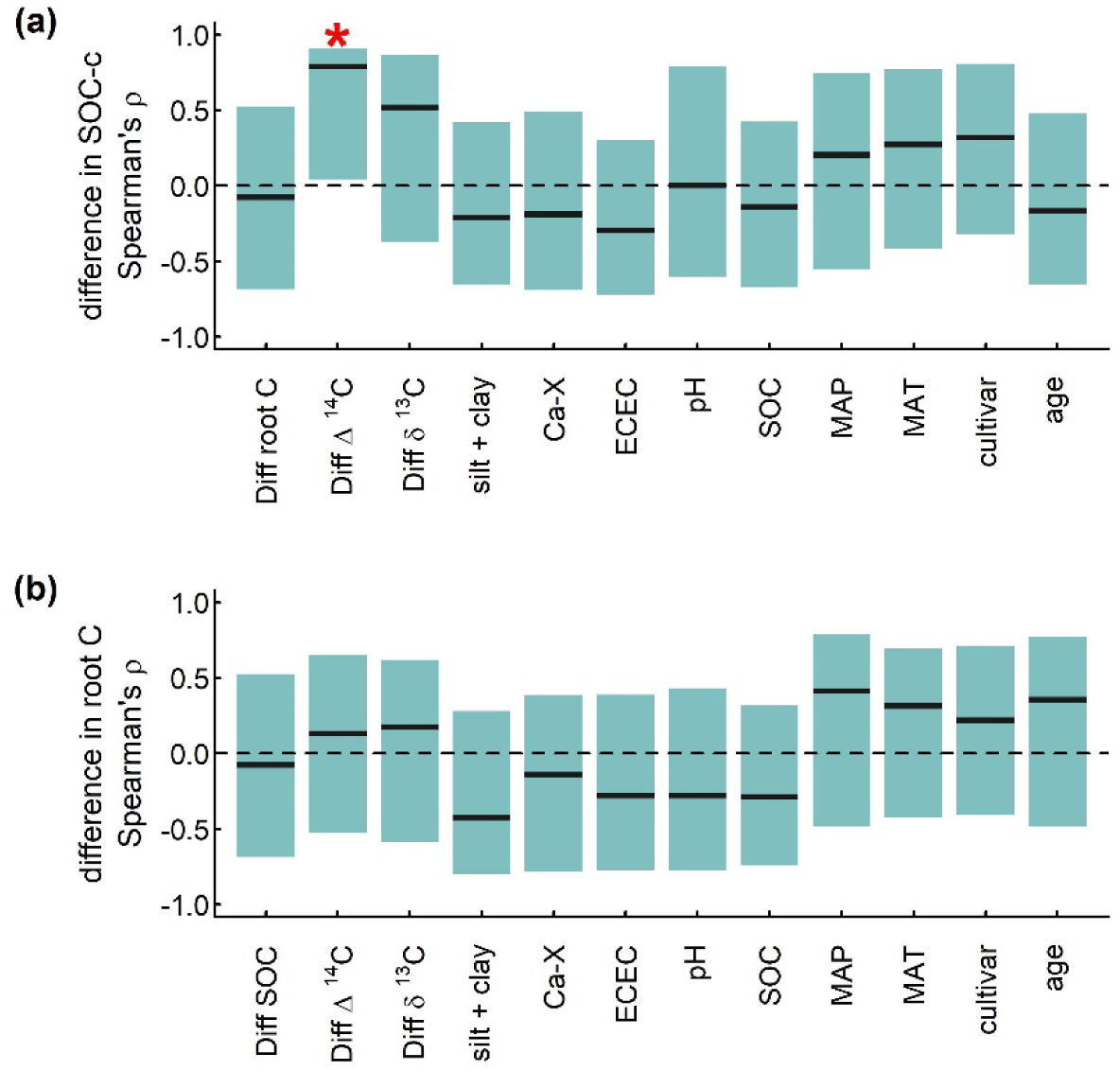
Correlation between differences in C pools between shallow-rooted annual crop fields and adjacent mature switchgrass and environmental variables at 12 sites across the central and eastern USA. Panels show Spearman’s ρ coefficient for paired correlations (n=12) with the difference (switchgrass – shallow-rooted annual crop) in texture-corrected C stock (a) and root C stock (b), computed after summarizing the data over the uppermost 0-1400 kg m^-2^ of mineral soil. Correlations were computed for each of the twelve variables listed on the horizontal axes. Black vertical lines show the observed across-site correlation statistic and shaded regions show bootstrapped 95% confidence intervals. Ca-X refers to exchangeable Ca^2+^, ECEC is the effective cation exchange capacity (charge-equivalent sum of BaCl_2_ extractable cations), and SOC represents the mean uncorrected SOC stock at each site. Switchgrass cultivar was coded as a binary variable so that upland cultivars = 1 and lowland cultivars = 2; hence positive correlation suggests a higher difference where lowland cultivars are present. A red asterisk indicates correlations for which the confidence interval does not include zero.

We further explored the correlation between the difference in SOC and the difference in Δ^14^C between vegetation types. We determined that this correlation was statistically significant even after accounting for multiple comparisons across all correlations tested (p = 0.03, Bonferroni adjusted one-sided permutation test). Examining this relationship more closely across the uppermost 0-1400 kg of soil, we found that the mean difference in SOC between switchgrass and shallow-rooted annual crops at a given site was linearly related to the mean difference in Δ^14^C between the two plant covers (Fig. 6a). We used a mixing model (Methods, Eq. 1) that used ^14^C to estimate the fraction of additional recently fixed SOC under switchgrass and plotted this value against the relative difference in SOC under the two plant covers [relative difference = (SOC switchgrass – SOC shallow-rooted annual crop)/SOC switchgrass]. These two relative difference values were linearly related and had comparable magnitudes (Fig. 6b). We compared the relative differences derived from ^14^C versus SOC and found that on average there was no statistically significant difference between the two metrics (mean difference = 0.09, paired t-test *p* = 0.25).

**Figure 6.**
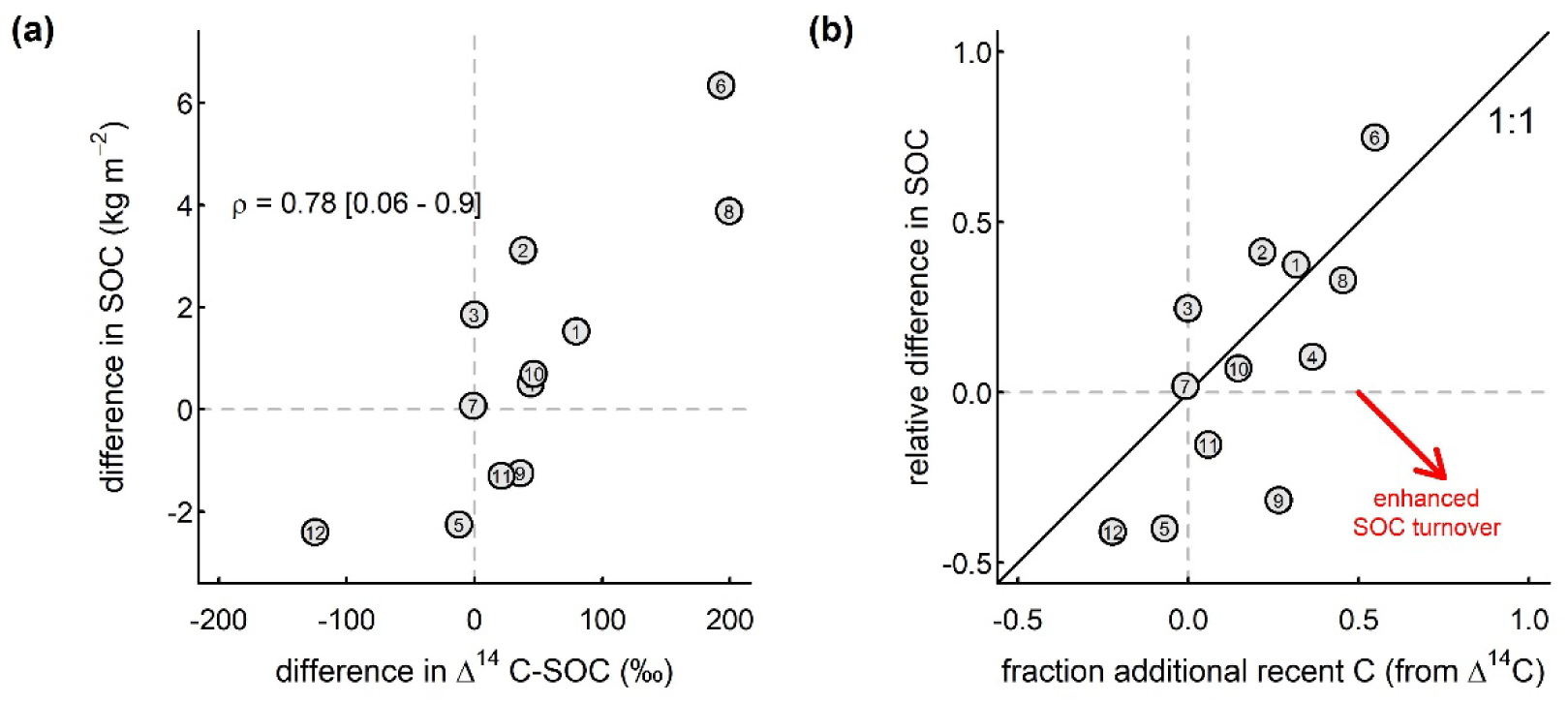
Effects of switchgrass cropping on SOC versus soil ^14^C at 12 sites across the central and eastern USA. Panel (a) shows the difference in Δ^14^C (switchgrass – shallow-rooted annual crop) versus the difference in SOC (switchgrass – shallow-rooted annual crop), correcting for texture variation within each site. Numbers correspond to the sites listed in Table 1. Spearman’s ρ calculated for these two variables equaled 0.78 (95% CI [0.06 to 0.9]). Panel (b) shows the relative difference in SOC (switchgrass – shallow-rooted annual crop)/(switchgrass) versus the fraction of additional recent C under switchgrass, estimated by applying a mixing model to the ^14^C data. The diagonal line represents a 1:1 relationship.

## 4 Discussion

### 4.1 Switchgrass had no clear effect on SOC but increased root C

We found little support for the hypothesis that replacing shallow-rooted annual crops with switchgrass increases SOC. SOC was higher on average under switchgrass in the surface 420 kg of soil (approximately 30 cm) after correcting for texture variability at each site. However, at the whole profile scale, differences in SOC were not statistically detectable, regardless of whether a texture correction was performed. Even though the majority of sites had more SOC under switchgrass, three sites had substantially less SOC under switchgrass (sites 5, 11, and 12), making the average effect highly uncertain. Drawing robust conclusions about SOC dynamics at depth requires a large number of replicate samples because the effects of management at depth are small relative to natural variability (Kravchenko and Robertson, 2011). Additionally, the dynamics of SOC recovery following perennial grassland restoration are slow: one study conducted near our site 10 estimated that full recovery of SOC would take more than 100 years (Matamala et al., 2008). Changes in SOC at this timescale, if they occurred, would be difficult to detect against background temporal variability.

While the average effect of switchgrass on SOC across sites was not statistically distinguishable from zero, a positive effect of switchgrass on root biomass C was clear. At the time of sampling, root biomass under switchgrass exceeded root biomass under shallow-rooted annual crops at virtually all locations and depths sampled. Furthermore, the depth at which 95% of roots were encountered was 62 cm greater under switchgrass than under shallow-rooted annual crops, indicating that at the time of sampling, switchgrass roots were not only more abundant but also more deeply distributed than annual crop roots. Accounting for root biomass C increased our overall estimate of belowground C under switchgrass by 0.6 kg C m^-2^. Given this contribution from roots, we have relatively high confidence that switchgrass increased overall belowground organic C storage (root C + SOC, Fig. 3d).

While it is clear that switchgrass cultivation increases root C stocks, our stock estimates are subject to a few caveats. First, we sampled under switchgrass crowns to characterize rooting distributions effectively. Sampling the gaps between individual switchgrass plants likely would have yielded lower estimates of root biomass C (Ma et al., 2000). Second, we sampled prior to flowering throughout the course of the summer growing season. While root biomass was likely higher during the summer growing season than during other times of year, we did not necessarily capture peak root biomass. Caveats aside, it is notable that the average amount of additional root C under switchgrass (0.6 kg C m^-2^) is comparable to the amount of additional SOC under switchgrass identified by a previous paired-plot study across the midwestern USA [0.77 kg C m^-2^ (Liebig et al., 2005)]. Additionally, perennial cropping systems maintain root stocks year round, and switchgrass has been shown to sustain substantial root biomass throughout all seasons (Dohleman et al., 2012). These observations suggest that root biomass might contribute significantly to belowground C storage if perennial plant cover and undisturbed soil are maintained over time.

### 4.2 C isotopes reflect new switchgrass C at depth

Carbon isotopes can reveal subtle time-integrated changes in the size or turnover of SOC that are not apparent from bulk SOC measurements (Balesdent and Mariotti, 1996; Baisden and Parfitt, 2007; Slessarev et al., 2020). The ^14^C and ^13^C signatures of the diverse soils that we sampled suggest these soils had experienced a range of historical C inputs and provide context for understanding the effects of switchgrass on SOC. For instance, ^14^C values varied substantially and reached extremely low values at depth, approaching −1000 ‰, which suggests that deep SOC at some sites was cycling at multi-millennial timescales. In many cases, these extremely low ^14^C values may have been caused by radiocarbon-dead petrogenic C (Keller and Bacon, 1998; Grant et al., 2023). We suspect that petrogenic organic C was abundant at several sites featuring glacially deposited sedimentary parent material (e.g., sites 8-12). The δ^13^C values that we observed at depth at these sites also reached extremely low values (< −30 ‰), consistent with the presence of petrogenic C (Longbottom and Hockaday, 2019). The diversity of δ^13^C values further suggests a mixed contribution of C_3_ and C_4_ vegetation to SOC across the sites. This pattern aligns with the fact that, in addition to switchgrass, which is C_4_, many of the sites would have hosted mixed C_3_ and C_4_ prairie grasses and C_3_ forbs pre-cultivation. Modern day agricultural inputs also include both C_3_ (e.g., soy, wheat) and C_4_ (e.g., corn) inputs.

Switchgrass tended to increase ^14^C values, shifting them towards the composition of the atmosphere, particularly at depth. Enrichment of ^14^C indicates that switchgrass transferred more newly fixed C to the subsoil than shallow-rooted annual crops. Applying a mixing model to the ^14^C data revealed that the relative increase in ^14^C at each site was comparable in magnitude to the relative difference in SOC (Fig. 6b). This finding suggests that increases in new C input under switchgrass is generally accompanied by a proportionate increase in the SOC stock: on average, newly fixed C was not simply replacing old C, but adding to existing SOC. Alternatively, newly added C could have stimulated decomposition of SOC (Kuzyakov, 2010). In this case, newly fixed C would have enhanced decomposition to the point that it mostly replaces the lost older C with little or no increase in total SOC. We did not see strong evidence that switchgrass enhanced decomposition across sites (Fig. 6b), although we cannot rule out this possibility at individual sites. For instance, at sites 9 and 11, we found greater relative shifts in ^14^C than in SOC, suggesting enhanced decomposition. On the other hand, in a parallel incubation study at these sites, we found that switchgrass enhanced respiration of newly fixed SOC rather than older SOC in topsoil (Min et al., 2025). Ultimately, the isotope mixing model results are best interpreted across all 12 sites given limited within-site replication.

Measurements of ^13^C also indicated that switchgrass was adding additional C through the surface 100 cm of soil. Unfortunately, the complex mix of plant inputs across our sites precluded applying a simple mixing model to relate the ^13^C data to bulk SOC dynamics. However, the patterns we observed qualitatively suggest that switchgrass enhanced C inputs below 30 cm.

### 4.3 No clear moderators of belowground C change

We found no clear evidence that any of the edaphic or environmental variables we measured moderated the effect of switchgrass on SOC. Indeed, we found no correlation between the effect of switchgrass on root C stocks and its effect on SOC stocks. We nonetheless consider a range of potential moderators that might be important for future studies. Of the twelve sites we sampled, four of the six sites where the effect of switchgrass on SOC was highest were more southerly, including two out of three of the sites in Oklahoma and both sites in North Carolina. These four sites featured several environmental factors in common: all hosted “lowland” switchgrass ecotypes (Kanlow or Alamo), all featured a relatively warm climate, and all had relatively coarse-textured soils with lower SOC stocks. Lowland switchgrass ecotypes generally produce more biomass than upland cultivars (Wullschleger et al., 2010); higher temperatures in the southern US would tend to increase evaporative demand and growing season length; and coarser textured soils tend to lower mean soil water content. Combined, these factors might encourage more root production in deeper soil layers and hence more SOC storage.

Contrary to one of our hypotheses, we found no clear role for soil chemical factors in moderating SOC accrual. Outside the context of this study, numerous lines of evidence point to the role of divalent cations—particularly Ca^2+^—in linking SOC to minerals in neutral to alkaline soils (Morris et al., 2007; O’Brien et al., 2015; Rowley et al., 2021; Shabtai et al., 2023) and even acidic soils (Rowley et al., 2023). Indeed, the correlation between *total* SOC stocks and exchangeable Ca in the uppermost ∼100 cm of soil across all sampled soil cores was relatively high (Spearman’s ρ = 0.66). It could be that the moderating effect of divalent cations on the response of SOC to land use is too weak to be detected across a limited number of sites. Alternatively, it could be that large-scale spatial correlations between SOC and divalent cations only emerge over longer timescales under natural conditions.

Interestingly, our analysis did not reveal any relationship between the effect of switchgrass on SOC and the duration since land management diverged in the two vegetation types, which ranged from 4 to 48 years. This result does not necessarily preclude a moderating effect of time on SOC accrual. Indeed, some relationship with time seems inevitable given that SOC cannot accrue instantaneously. The sites that we sampled varied widely along environmental axes and were not controlled for tillage or fertilization practices; hence, these sites are not suitable for constructing a chronosequence of SOC accrual under switchgrass. It is nonetheless striking that sites with land use contrasts in the range of 4 to 48 years showed little evidence of progressive SOC accrual over time. Clearly, the trajectory of SOC change following switchgrass cultivation is slow enough that the role of time can be overwhelmed by other factors. The dynamics of SOC recovery after conversion of cropland to perennial grassland are also not necessarily monotonically increasing. Cessation of tillage and associated aboveground litter burial can cause SOC to decrease at depth during the initial stages of grassland restoration (O’Brien et al., 2010). Describing the details of these dynamics would be best achieved with a site-level time series of soil sampling rather than regression across multiple sites.

One factor that we did not quantify or consider in our analysis was plant community diversity, which might also moderate the effect of deep-rooted perennials on SOC. At site 9, one study found that more diverse grassland communities support higher fine root production (Sprunger et al., 2017). At site 11, more diverse cropping systems scored higher on a range of soil health metrics (Sanford et al., 2021), and relatively diverse prairie plots retained more SOC and supported high microbial carbon use efficiency than lower diversity cropping systems (Liang et al., 2016; Rui et al., 2022; Dietz et al., 2024). The switchgrass fields that we sampled varied in diversity from pure monocultures to stands that included some forbs (e.g., invaded by *Solidago* spp.). Future regional surveys like ours might consider diversity quantitatively.

### 4.4 Implications for cropland SOC management and climate

Perennial grasses might be introduced to agricultural systems by planting perennial food crops (e.g., intermediate wheatgrass; (Culman et al., 2013)), planting biomass crops on marginal lands, or by restoring grasslands in low-yielding croplands (Cai et al., 2011; Kell, 2012; Lynch and Wojciechowski, 2015; Conant et al., 2017; Schulte et al., 2017; Yang and Tilman, 2020). The effect of these strategies on atmospheric CO_2_ levels depends on complex economic factors related to land use. For instance, establishing lower-yielding perennial grains, biomass crops, or restored grassland on active cropland may affect commodity prices, driving expansion of agriculture and SOC loss elsewhere (Plevin et al., 2010). The SOC that accrues under perennial grasses is also vulnerable to loss if future economic conditions incentivize re-cultivation of restored grassland (Moinet et al., 2023). Here we focus on short-term local changes in SOC storage while acknowledging that the effect of land use change on Earth’s atmosphere ultimately depends on these economic factors.

Taking this short-term perspective, our study adds to the growing body of evidence that cropland management can affect belowground C at depths > 30 cm. While we did not find evidence of consistent SOC change at depth, we did find evidence that total belowground carbon stocks (i.e., SOC plus root carbon stock) to a depth of 100 cm were positively impacted by cultivation of a deep-rooted perennial. Our isotopic measurements also indicated that switchgrass was adding new C at depth. We found root biomass under switchgrass at depths well over 100 cm (Fig. 2b), suggesting that planting perennial grasses can affect biogeochemical processes below the depths commonly considered during agronomic sampling.

We also found that increases in root C can be significant. This finding supports the idea that root biomass should be explicitly included when quantifying C dynamics in life-cycle analyses of agricultural systems (Gelfand et al., 2020). While root biomass is readily decomposed, SOC and aboveground biomass are also non-durable reservoirs for storing C. Climate benefits from increasing root C, SOC, or aboveground biomass are thus all ultimately dependent on our capacity to maintain beneficial management practices long enough for more durable decarbonization to unfold (Mayer et al., 2025). Promoting C storage in non-durable ecological pools like root biomass is no substitute for eliminating fossil fuel emissions (Schenuit et al., 2023). However, our findings show that restoring and maintaining deep rooted perennial grasses can help to counteract belowground C loss due to land use, benefiting Earth’s climate.

Perennial cropping systems also produce more stable yields and score better on soil health indices than annual cropping systems (Sanford et al., 2021), while eliminating the need for tillage and associated erosion. Taking this holistic perspective, there are many potential environmental benefits associated with deep-rooted perennials beyond direct climate effects.

## Acknowledgements

We thank the following individuals for facilitating site access and providing logistical support during field work: Brandon Carr (Bud Smith Plant Materials Center, Texas), Christine Hawkes (NC State University), David Lang, Mississippi State University, under MIS-CRIS 161090, Wally Levernier (Fermi National Accelerator Laboratory), and Michael Ricketts (Argonne National Laboratory). Christy Ramon (LLNL) assisted with laboratory sample prep. This work was supported by Lawrence Livermore National Laboratory (LLNL)’s Lab Directed Research and Development program (#19-ERD-010); a DOE Early Career Research Program award to EN (SCW1711); and the Center for Advanced Bioenergy and Bioproducts Innovation Contract DE-AC02-05CH11231, Award Number DE-SC0018420. Work at LLNL was conducted under the auspices of the Department of Energy Contract DE-AC52-07NA27344. Argonne National Laboratory’s contribution was supported by the U.S. Department of Energy, Office of Science, Biological and Environmental Research Program under contract DE-AC02-06CH11357.

## Data Availability

The data described in this study are publicly available as .csv files on Zenodo at doi:10.5281/zenodo.16620529. R code used to perform bootstrapping analyses is included as Supplementary Information.

## References

Abatzoglou, J.T., 2013. Development of gridded surface meteorological data for ecological applications and modelling. International Journal of Climatology 33, 121–131. doi:10.1002/joc.3413

Adkins, J., Jastrow, J.D., Morris, G.P., De Graaff, M.-A., 2019. Effects of fertilization, plant species, and intra-specific diversity on soil carbon and nitrogen in biofuel cropping systems after five growing seasons. Biomass and Bioenergy 130, 105393. doi:10.1016/j.biombioe.2019.105393

Assad, E.D., Pinto, H.S., Martins, S.C., Groppo, J.D., Salgado, P.R., Evangelista, B., Vasconcellos, E., Sano, E.E., Pavão, E., Luna, R., Camargo, P.B., Martinelli, L.A., 2013. Changes in soil carbon stocks in Brazil due to land use: paired site comparisons and a regional pasture soil survey. Biogeosciences 10, 6141–6160. doi:10.5194/bg-10-6141-2013

Austin, E.E., Wickings, K., McDaniel, M.D., Robertson, G.P., Grandy, A.S., 2017. Cover crop root contributions to soil carbon in a no-till corn bioenergy cropping system. GCB Bioenergy 9, 1252–1263. doi:10.1111/gcbb.12428

Baisden, W.T., Parfitt, R.L., 2007. Bomb 14C enrichment indicates decadal C pool in deep soil? Biogeochemistry 85, 59–68. doi:10.1007/s10533-007-9101-7

Balesdent, J., Basile-Doelsch, I., Chadoeuf, J., Cornu, S., Derrien, D., Fekiacova, Z., Hatté, C., 2018. Atmosphere–soil carbon transfer as a function of soil depth. Nature 559, 599–602. doi:10.1038/s41586-018-0328-3

Balesdent, J., Mariotti, A., 1996. Measurment of turnover of soil organic matter using 13C natural abundance, in: Mass Spectrometry of Soils. Marcel Dekker, New York.

Becker, A.E., Horowitz, L.S., Ruark, M.D., Jackson, R.D., 2022. Surface-soil carbon stocks greater under well-managed grazed pasture than row crops. Soil Science Society of America Journal 86, 758–768. doi:10.1002/saj2.20388

Beillouin, D., Corbeels, M., Demenois, J., Berre, D., Boyer, A., Fallot, A., Feder, F., Cardinael, R., 2023. A global meta-analysis of soil organic carbon in the Anthropocene. Nature Communications 14, 3700. doi:10.1038/s41467-023-39338-z

Biondini, M.E., Patton, B.D., Nyren, P.E., 1998. Grazing Intensity and Ecosystem Processes in a Northern Mixed-Grass Prairie, USA. Ecological Applications 8, 469–479. doi:10.1890/1051-0761(1998)008[0469:GIAEPI]2.0.CO;2

Bossio, D.A., Cook-Patton, S.C., Ellis, P.W., Fargione, J., Sanderman, J., Smith, P., Wood, S., Zomer, R.J., Von Unger, M., Emmer, I.M., Griscom, B.W., 2020. The role of soil carbon in natural climate solutions. Nature Sustainability 3, 391–398. doi:10.1038/s41893-020-0491-z

Burt, R., 2014. Soil Survey Laboratory Methods Manual (No 42. Version 5).

Cai, X., Zhang, X., Wang, D., 2011. Land Availability for Biofuel Production. Environmental Science & Technology 45, 334–339. doi:10.1021/es103338e

Conant, R.T., Cerri, C.E.P., Osborne, B.B., Paustian, K., 2017. Grassland management impacts on soil carbon stocks: a new synthesis. Ecological Applications 27, 662–668. doi:10.1002/eap.1473

Culman, S.W., Snapp, S.S., Ollenburger, M., Basso, B., DeHaan, L.R., 2013. Soil and Water Quality Rapidly Responds to the Perennial Grain Kernza Wheatgrass. Agronomy Journal 105, 735–744. doi:10.2134/agronj2012.0273

Das, S., Teuffer, K., Stoof, C.R., Walter, M.F., Walter, M.T., Steenhuis, T.S., Richards, B.K., 2018. Perennial Grass Bioenergy Cropping on Wet Marginal Land: Impacts on Soil Properties, Soil Organic Carbon, and Biomass During Initial Establishment. BioEnergy Research 11, 262–276. doi:10.1007/s12155-018-9893-4

Davison, A.C., Hinkley, D.V., 1997. Bootstrap methods and their application. Cambridge University Press, Cambridge ; New York, NY, USA.

Dietz, C.L., Jackson, R.D., Ruark, M.D., Sanford, G.R., 2024. Soil carbon maintained by perennial grasslands over 30 years but lost in field crop systems in a temperate Mollisol. Communications Earth & Environment 5, 360. doi:10.1038/s43247-024-01500-w

Dohleman, F.G., Heaton, E.A., Arundale, R.A., Long, S.P., 2012. Seasonal dynamics of above- and below-ground biomass and nitrogen partitioning in *M iscanthus* × *giganteus* and *P anicum virgatum* across three growing seasons. GCB Bioenergy 4, 534–544. doi:10.1111/j.1757-1707.2011.01153.x

Fontaine, S., Barot, S., Barré, P., Bdioui, N., Mary, B., Rumpel, C., 2007. Stability of organic carbon in deep soil layers controlled by fresh carbon supply. Nature 450, 277–280. doi:10.1038/nature06275

Gelfand, I., Hamilton, S.K., Kravchenko, A.N., Jackson, R.D., Thelen, K.D., Robertson, G.P., 2020. Empirical Evidence for the Potential Climate Benefits of Decarbonizing Light Vehicle Transport in the U.S. with Bioenergy from Purpose-Grown Biomass with and without BECCS. Environmental Science & Technology 54, 2961–2974. doi:10.1021/acs.est.9b07019

Gifford, R.M., Roderick, M.L., 2003. Soil carbon stocks and bulk density: spatial or cumulative mass coordinates as a basis of expression? Global Change Biology 9, 1507–1514. doi:10.1046/j.1365-2486.2003.00677.x

Grant, K.E., Hilton, R.G., Galy, V.V., 2023. Global patterns of radiocarbon depletion in subsoil linked to rock-derived organic carbon. Geochemical Perspectives Letters 25, 36–40. doi:10.7185/geochemlet.2312

Harris, D., Horwáth, W.R., Van Kessel, C., 2001. Acid fumigation of soils to remove carbonates prior to total organic carbon or CARBON-13 isotopic analysis. Soil Science Society of America Journal 65, 1853–1856. doi:10.2136/sssaj2001.1853

Hauser, E., Sullivan, P.L., Flores, A.N., Hirmas, D., Billings, S.A., 2022. Global-Scale Shifts in Rooting Depths Due To Anthropocene Land Cover Changes Pose Unexamined Consequences for Critical Zone Functioning. Earth’s Future 10, e2022EF002897. doi:10.1029/2022EF002897

Helal, H.M., Sauerbeck, D., 1986. Effect of plant roots on carbon metabolism of soil microbial biomass. Zeitschrift Für Pflanzenernährung Und Bodenkunde 149, 181–188. doi:10.1002/jpln.19861490205

Hicks Pries, C.E., Ryals, R., Zhu, B., Min, K., Cooper, A., Goldsmith, S., Pett-Ridge, J., Torn, M., Berhe, A.A., 2023. The Deep Soil Organic Carbon Response to Global Change. Annual Review of Ecology, Evolution, and Systematics 54, 375–401. doi:10.1146/annurev-ecolsys-102320-085332

Hirte, J., Leifeld, J., Abiven, S., Oberholzer, H.-R., Hammelehle, A., Mayer, J., 2017. Overestimation of Crop Root Biomass in Field Experiments Due to Extraneous Organic Matter. Frontiers in Plant Science 8. doi:10.3389/fpls.2017.00284

Hong, S., Yin, G., Piao, S., Dybzinski, R., Cong, N., Li, X., Wang, K., Peñuelas, J., Zeng, H., Chen, A., 2020. Divergent responses of soil organic carbon to afforestation. Nature Sustainability 3, 694–700. doi:10.1038/s41893-020-0557-y

Huo, C., Luo, Y., Cheng, W., 2017. Rhizosphere priming effect: A meta-analysis. Soil Biology and Biochemistry 111, 78–84. doi:10.1016/j.soilbio.2017.04.003

Kell, D.B., 2012. Large-scale sequestration of atmospheric carbon via plant roots in natural and agricultural ecosystems: why and how. Philosophical Transactions of the Royal Society B: Biological Sciences 367, 1589–1597. doi:10.1098/rstb.2011.0244

Keller, C.K., Bacon, D.H., 1998. Soil respiration and georespiration distinguished by transport analyses of vadose CO2,13 CO2, and14 CO2. Global Biogeochemical Cycles 12, 361–372. doi:10.1029/98GB00742

Kelly-Slatten, M.J., Stewart, C.E., Tfaily, M.M., Jastrow, J.D., Sasso, A., De Graaff, M., 2023. Root traits of perennial C4 grasses contribute to cultivar variations in soil chemistry and species patterns in particulate and mineral-associated carbon pool formation. GCB Bioenergy 15, 613–629. doi:10.1111/gcbb.13041

King, A.E., Amsili, J.P., Córdova, S.C., Culman, S., Fonte, S.J., Kotcon, J., Liebig, M., Masters, M.D., McVay, K., Olk, D.C., Schipanski, M., Schneider, S.K., Stewart, C.E., Cotrufo, M.F., 2023. A soil matrix capacity index to predict mineral-associated but not particulate organic carbon across a range of climate and soil pH. Biogeochemistry 165, 1–14. doi:10.1007/s10533-023-01066-3

King, A.E., Blesh, J., 2018. Crop rotations for increased soil carbon: perenniality as a guiding principle. Ecological Applications 28, 249–261. doi:10.1002/eap.1648

Kitchen, D.J., Blair, J.M., Callaham, M.A., 2009. Annual fire and mowing alter biomass, depth distribution, and C and N content of roots and soil in tallgrass prairie. Plant and Soil 323, 235–247. doi:10.1007/s11104-009-9931-2

Kravchenko, A.N., Robertson, G.P., 2011. Whole-Profile Soil Carbon Stocks: The Danger of Assuming Too Much from Analyses of Too Little. Soil Science Society of America Journal 75, 235–240. doi:10.2136/sssaj2010.0076

Kropko, J., Harden, J., 2020. coxed: Duration-Based Quantities of Interest for the Cox Proportional Hazards. R package version 0.3.3.

Kuzyakov, Y., 2010. Priming effects: Interactions between living and dead organic matter. Soil Biology and Biochemistry 42, 1363–1371. doi:10.1016/j.soilbio.2010.04.003

Lai, L., Kumar, S., Osborne, S., Owens, V.N., 2018. Switchgrass impact on selected soil parameters, including soil organic carbon, within six years of establishment. CATENA 163, 288–296. doi:10.1016/j.catena.2017.12.030

Lark, R.M., Bellamy, P.H., Kirk, G.J.D., 2006. Baseline values and change in the soil, and implications for monitoring. European Journal of Soil Science 57, 916–921. doi:10.1111/j.1365-2389.2006.00875.x

Ledo, A., Smith, P., Zerihun, A., Whitaker, J., Vicente-Vicente, J.L., Qin, Z., McNamara, N.P., Zinn, Y.L., Llorente, M., Liebig, M., Kuhnert, M., Dondini, M., Don, A., Diaz-Pines, E., Datta, A., Bakka, H., Aguilera, E., Hillier, J., 2020. Changes in soil organic carbon under perennial crops. Global Change Biology gcb.15120. doi:10.1111/gcb.15120

Liang, C., Kao-Kniffin, J., Sanford, G.R., Wickings, K., Balser, T.C., Jackson, R.D., 2016. Microorganisms and their residues under restored perennial grassland communities of varying diversity. Soil Biology and Biochemistry 103, 192–200. doi:10.1016/j.soilbio.2016.08.002

Liebig, M.A., Johnson, H.A., Hanson, J.D., Frank, A.B., 2005. Soil carbon under switchgrass stands and cultivated cropland. Biomass and Bioenergy 28, 347–354. doi:10.1016/j.biombioe.2004.11.004

Longbottom, T.L., Hockaday, W.C., 2019. Molecular and isotopic composition of modern soils derived from kerogen-rich bedrock and implications for the global C cycle. Biogeochemistry 143, 239–255. doi:10.1007/s10533-019-00559-4

Lynch, J.P., Wojciechowski, T., 2015. Opportunities and challenges in the subsoil: pathways to deeper rooted crops. Journal of Experimental Botany 66, 2199–2210. doi:10.1093/jxb/eru508

Ma, Z., Wood, C.W., Bransby, D.I., 2000. Soil management impacts on soil carbon sequestration by switchgrass. Biomass and Bioenergy 18, 469–477. doi:10.1016/S0961-9534(00)00013-1

Matamala, R., Jastrow, J.D., Miller, R.M., Garten, C.T., 2008. Temporal Changes in C and N Stocks of Restored Prairie: Implications for C Sequestraion Strategies. Ecological Applications 18, 1470–1488. doi:10.1890/07-1609.1

Mayer, A., Dumortier, J., Hausfather, Z., Pett-Ridge, J., Slessarev, E., 2025. Cost of cooling: The value of reversible carbon storage in a zero-emissions world. doi:10.70212/cdrxiv.2025348.v1

Min, K., Nuccio, E., Slessarev, E., Kan, M., McFarlane, K.J., Oerter, E., Jurusik, A., Sanford, G., Thelen, K.D., Pett-Ridge, J., Berhe, A.A., 2025. Deep-rooted perennials alter microbial respiration and chemical composition of carbon in density fractions along soil depth profiles. Geoderma 455, 117202. doi:10.1016/j.geoderma.2025.117202

Min, K., Slessarev, E., Kan, M., McFarlane, K., Oerter, E., Pett-Ridge, J., Nuccio, E., Berhe, A.A., 2021. Active microbial biomass decreases, but microbial growth potential remains similar across soil depth profiles under deeply-vs. shallow-rooted plants. Soil Biology and Biochemistry 162, 108401. doi:10.1016/j.soilbio.2021.108401

Moinet, G.Y.K., Hijbeek, R., Van Vuuren, D.P., Giller, K.E., 2023. Carbon for soils, not soils for carbon. Global Change Biology 29, 2384–2398. doi:10.1111/gcb.16570

Moore, E.B., De, M., Nunes, M.R., Saha, D., Jin, V., Li, L., Johnson, J.M.F., Karlen, D.L., McDaniel, M.D., 2025. Connections between roots and soil health across agriculture management practices. Plant and Soil. doi:10.1007/s11104-025-07367-w

Morris, S.J., Bohm, S., Haile-Mariam, S., Paul, E.A., 2007. Evaluation of carbon accrual in afforested agricultural soils. Global Change Biology 13, 1145–1156. doi:10.1111/j.1365-2486.2007.01359.x

Mosier, S., Apfelbaum, S., Byck, P., Calderon, F., Teague, R., Thompson, R., Cotrufo, M.F., 2021. Adaptive multi-paddock grazing enhances soil carbon and nitrogen stocks and stabilization through mineral association in southeastern U.S. grazing lands. Journal of Environmental Management 288, 112409. doi:10.1016/j.jenvman.2021.112409

O’Brien, S.L., Jastrow, J.D., Grimley, D.A., Gonzalez-Meler, M.A., 2015. Edaphic controls on soil organic carbon stocks in restored grasslands. Geoderma 251–252, 117–123. doi:10.1016/j.geoderma.2015.03.023

O’Brien, S.L., Jastrow, J.D., Grimley, D.A., Gonzalez-Meler, M.A., 2010. Moisture and vegetation controls on decadal-scale accrual of soil organic carbon and total nitrogen in restored grasslands. Global Change Biology 16, 2573–2588. doi:10.1111/j.1365-2486.2009.02114.x

Ofiti, N.O.E., Zosso, C.U., Soong, J.L., Solly, E.F., Torn, M.S., Wiesenberg, G.L.B., Schmidt, M.W.I., 2021. Warming promotes loss of subsoil carbon through accelerated degradation of plant-derived organic matter. Soil Biology and Biochemistry 156, 108185. doi:10.1016/j.soilbio.2021.108185

Ogle, S.M., Breidt, F.J., Paustian, K., 2005. Agricultural management impacts on soil organic carbon storage under moist and dry climatic conditions of temperate and tropical regions. Biogeochemistry 72, 87–121. doi:10.1007/s10533-004-0360-2

Olson, K.R., Al-Kaisi, M., Lal, R., Lowery, B., 2014. Examining the paired comparison method approach for determining soil organic carbon sequestration rates. Journal of Soil and Water Conservation 69, 193A–197A. doi:10.2489/jswc.69.6.193A

Padarian, J., Stockmann, U., Minasny, B., McBratney, A.B., 2022. Monitoring changes in global soil organic carbon stocks from space. Remote Sensing of Environment 281, 113260. doi:10.1016/j.rse.2022.113260

Peixoto, L., Olesen, J.E., Elsgaard, L., Enggrob, K.L., Banfield, C.C., Dippold, M.A., Nicolaisen, M.H., Bak, F., Zang, H., Dresbøll, D.B., Thorup-Kristensen, K., Rasmussen, J., 2022. Deep-rooted perennial crops differ in capacity to stabilize C inputs in deep soil layers. Scientific Reports 12, 5952. doi:10.1038/s41598-022-09737-1

Plevin, R.J., Jones, A.D., Torn, M.S., Gibbs, H.K., 2010. Greenhouse Gas Emissions from Biofuels’ Indirect Land Use Change Are Uncertain but May Be Much Greater than Previously Estimated. Environmental Science & Technology 44, 8015–8021. doi:10.1021/es101946t

Poeplau, C., Don, A., Schneider, F., 2021. Roots are key to increasing the mean residence time of organic carbon entering temperate agricultural soils. Global Change Biology 27, 4921– 4934. doi:10.1111/gcb.15787

Pribyl, D.W., 2010. A critical review of the conventional SOC to SOM conversion factor. Geoderma 156, 75–83. doi:10.1016/j.geoderma.2010.02.003

Rasmussen, C., Heckman, K., Wieder, W.R., Keiluweit, M., Lawrence, C.R., Berhe, A.A., Blankinship, J.C., Crow, S.E., Druhan, J.L., Hicks Pries, C.E., Marin-Spiotta, E., Plante, A.F., Schädel, C., Schimel, J.P., Sierra, C.A., Thompson, A., Wagai, R., 2018. Beyond clay: towards an improved set of variables for predicting soil organic matter content. Biogeochemistry 137, 297–306. doi:10.1007/s10533-018-0424-3

Rasse, D.P., Rumpel, C., Dignac, M.-F., 2005. Is soil carbon mostly root carbon? Mechanisms for a specific stabilisation. Plant and Soil 269, 341–356. doi:10.1007/s11104-004-0907-y

Rovira, P., Sauras, T., Salgado, J., Merino, A., 2015. Towards sound comparisons of soil carbon stocks: A proposal based on the cumulative coordinates approach. CATENA 133, 420–431. doi:10.1016/j.catena.2015.05.020

Rowley, M.C., Grand, S., Spangenberg, J.E., Verrecchia, E.P., 2021. Evidence linking calcium to increased organo-mineral association in soils. Biogeochemistry 153, 223–241. doi:10.1007/s10533-021-00779-7

Rowley, M.C., Grand, S., Verrecchia, É.P., 2018. Calcium-mediated stabilisation of soil organic carbon. Biogeochemistry 137, 27–49. doi:10.1007/s10533-017-0410-1

Rowley, M.C., Nico, P.S., Bone, S.E., Marcus, M.A., Pegoraro, E.F., Castanha, C., Kang, K., Bhattacharyya, A., Torn, M.S., Peña, J., 2023. Association between soil organic carbon and calcium in acidic grassland soils from Point Reyes National Seashore, CA. Biogeochemistry 165, 91–111. doi:10.1007/s10533-023-01059-2

Rui, Y., Jackson, R.D., Cotrufo, M.F., Sanford, G.R., Spiesman, B.J., Deiss, L., Culman, S.W., Liang, C., Ruark, M.D., 2022. Persistent soil carbon enhanced in Mollisols by well-managed grasslands but not annual grain or dairy forage cropping systems. Proceedings of the National Academy of Sciences 119, e2118931119. doi:10.1073/pnas.2118931119

Sanderman, J., Baldock, J.A., 2010. Accounting for soil carbon sequestration in national inventories: a soil scientist’s perspective. Environmental Research Letters 5, 034003. doi:10.1088/1748-9326/5/3/034003

Sanderman, J., Hengl, T., Fiske, G.J., 2017. Soil carbon debt of 12,000 years of human land use 113, 9575–9580. doi:doi/10.1073/pnas.1706103114

Sanford, G.R., Jackson, R.D., Booth, E.G., Hedtcke, J.L., Picasso, V., 2021. Perenniality and diversity drive output stability and resilience in a 26-year cropping systems experiment. Field Crops Research 263, 108071. doi:10.1016/j.fcr.2021.108071

Sanford, G.R., Posner, J.L., Jackson, R.D., Kucharik, C.J., Hedtcke, J.L., Lin, T.-L., 2012. Soil carbon lost from Mollisols of the North Central U.S.A. with 20 years of agricultural best management practices. Agriculture, Ecosystems & Environment 162, 68–76. doi:10.1016/j.agee.2012.08.011

Schenk, H.J., Jackson, R.B., 2005. Mapping the global distribution of deep roots in relation to climate and soil characteristics. Geoderma 126, 129–140. doi:10.1016/j.geoderma.2004.11.018

Schenuit, F., Gidden, M.J., Boettcher, M., Brutschin, E., Fyson, C., Gasser, T., Geden, O., Lamb, W.F., Mace, M.J., Minx, J., Riahi, K., 2023. Secure robust carbon dioxide removal policy through credible certification. Communications Earth & Environment 4, 349. doi:10.1038/s43247-023-01014-x

Schulte, L.A., Niemi, J., Helmers, M.J., Liebman, M., Arbuckle, J.G., James, D.E., Kolka, R.K., O’Neal, M.E., Tomer, M.D., Tyndall, J.C., Asbjornsen, H., Drobney, P., Neal, J., Van Ryswyk, G., Witte, C., 2017. Prairie strips improve biodiversity and the delivery of multiple ecosystem services from corn–soybean croplands. Proceedings of the National Academy of Sciences 114, 11247–11252. doi:10.1073/pnas.1620229114

Shabtai, I.A., Wilhelm, R.C., Schweizer, S.A., Höschen, C., Buckley, D.H., Lehmann, J., 2023. Calcium promotes persistent soil organic matter by altering microbial transformation of plant litter. Nature Communications 14, 6609. doi:10.1038/s41467-023-42291-6

Sierra, C.A., Müller, M., Trumbore, S.E., 2012. Models of soil organic matter decomposition: the SoilR package, version 1.0. Geoscientific Model Development 5, 1045–1060. doi:10.5194/gmd-5-1045-2012

Skadell, L.E., Schneider, F., Gocke, M.I., Guigue, J., Amelung, W., Bauke, S.L., Hobley, E.U., Barkusky, D., Honermeier, B., Kögel-Knabner, I., Schmidhalter, U., Schweitzer, K., Seidel, S.J., Siebert, S., Sommer, M., Vaziritabar, Y., Don, A., 2023. Twenty percent of agricultural management effects on organic carbon stocks occur in subsoils – Results of ten long-term experiments. Agriculture, Ecosystems & Environment 356, 108619. doi:10.1016/j.agee.2023.108619

Slessarev, E.W., Chadwick, O.A., Sokol, N.W., Nuccio, E.E., Pett-Ridge, J., 2022a. Rock weathering controls the potential for soil carbon storage at a continental scale. Biogeochemistry 157, 1–13. doi:10.1007/s10533-021-00859-8

Slessarev, E.W., Mayer, A., Kelly, C., Georgiou, K., Pett-Ridge, J., Nuccio, E.E., 2022b. Initial soil organic carbon stocks govern changes in soil carbon: reality or artifact? Global Change Biology gcb.16491. doi:10.1111/gcb.16491

Slessarev, E.W., Nuccio, E.E., McFarlane, K.J., Ramon, C., Saha, M., Firestone, M.K., Pett-Ridge, J., 2020. Quantifying the effects of switchgrass (*Panicum virgatum*) on deep organic C stocks using natural abundance ^14^ C in three marginal soils. GCB Bioenergy gcbb.12729. doi:10.1111/gcbb.12729

Sokol, N.W., Kuebbing, Sara.E., Karlsen-Ayala, E., Bradford, M.A., 2019. Evidence for the primacy of living root inputs, not root or shoot litter, in forming soil organic carbon. New Phytologist 221, 233–246. doi:10.1111/nph.15361

Soong, J.L., Castanha, C., Hicks Pries, C.E., Ofiti, N., Porras, R.C., Riley, W.J., Schmidt, M.W.I., Torn, M.S., 2021. Five years of whole-soil warming led to loss of subsoil carbon stocks and increased CO _2_ efflux. Science Advances 7, eabd1343. doi:10.1126/sciadv.abd1343

Sprunger, C.D., Oates, L.G., Jackson, R.D., Robertson, G.P., 2017. Plant community composition influences fine root production and biomass allocation in perennial bioenergy cropping systems of the upper Midwest, USA. Biomass and Bioenergy 105, 248–258. doi:10.1016/j.biombioe.2017.07.007

Sprunger, C.D., Robertson, P.G., 2018. Early accumulation of active fraction soil carbon in newly established cellulosic biofuel systems. Geoderma 318, 42–51. doi:10.1016/j.geoderma.2017.11.040

Stoner, S.W., Hoyt, A.M., Trumbore, S., Sierra, C.A., Schrumpf, M., Doetterl, S., Baisden, W.T., Schipper, L.A., 2021. Soil organic matter turnover rates increase to match increased inputs in grazed grasslands. Biogeochemistry 156, 145–160. doi:10.1007/s10533-021-00838-z

Sulman, B.N., Harden, J., He, Y., Treat, C., Koven, C., Mishra, U., O’Donnell, J.A., Nave, L.E., 2020. Land Use and Land Cover Affect the Depth Distribution of Soil Carbon: Insights From a Large Database of Soil Profiles. Frontiers in Environmental Science 8, 146. doi:10.3389/fenvs.2020.00146

VandenBygaart, A.J., Bremer, E., McConkey, B.G., Ellert, B.H., Janzen, H.H., Angers, D.A., Carter, M.R., Drury, C.F., Lafond, G.P., McKenzie, R.H., 2011. Impact of Sampling Depth on Differences in Soil Carbon Stocks in Long-Term Agroecosystem Experiments. Soil Science Society of America Journal 75, 226–234. doi:10.2136/sssaj2010.0099

Von Fromm, S.F., Hoyt, A.M., Lange, M., Acquah, G.E., Aynekulu, E., Berhe, A.A., Haefele,S.M., McGrath, S.P., Shepherd, K.D., Sila, A.M., Six, J., Towett, E.K., Trumbore, S.E., Vågen, T.-G., Weullow, E., Winowiecki, L.A., Doetterl, S., 2021. Continental-scale controls on soil organic carbon across sub-Saharan Africa. SOIL 7, 305–332. doi:10.5194/soil-7-305-2021

Von Haden, A.C., Dornbush, M.E., 2017. Ecosystem carbon pools, fluxes, and balances within mature tallgrass prairie restorations. Restoration Ecology 25, 549–558. doi:10.1111/rec.12461

Von Haden, A.C., Yang, W.H., DeLucia, E.H., 2020. Soils’ dirty little secret: Depth-based comparisons can be inadequate for quantifying changes in soil organic carbon and other mineral soil properties. Global Change Biology 26, 3759–3770. doi:10.1111/gcb.15124

Weaver, J.E., 1919. The Ecological Relations of Roots.

Wendt, J.W., Hauser, S., 2013. An equivalent soil mass procedure for monitoring soil organic carbon in multiple soil layers. European Journal of Soil Science 64, 58–65. doi:10.1111/ejss.12002

Wilsey, B.J., Polley, W.H., 2006. Aboveground productivity and root–shoot allocation differ between native and introduced grass species. Oecologia 150, 300–309. doi:10.1007/s00442-006-0515-z

Wullschleger, S.D., Davis, E.B., Borsuk, M.E., Gunderson, C.A., Lynd, L.R., 2010. Biomass Production in Switchgrass across the United States: Database Description and Determinants of Yield. Agronomy Journal 102, 1158–1168. doi:10.2134/agronj2010.0087

Yang, Y., Tilman, D., 2020. Soil and root carbon storage is key to climate benefits of bioenergy crops. Biofuel Research Journal 7, 1143–1148. doi:10.18331/BRJ2020.7.2.2

