## Supplemental figures 1-4 for "Assessing the Potential of Switchgrass (*Panicum virgatum* L.) for Storing Carbon Belowground: Insights from a Multi-Site US Study"

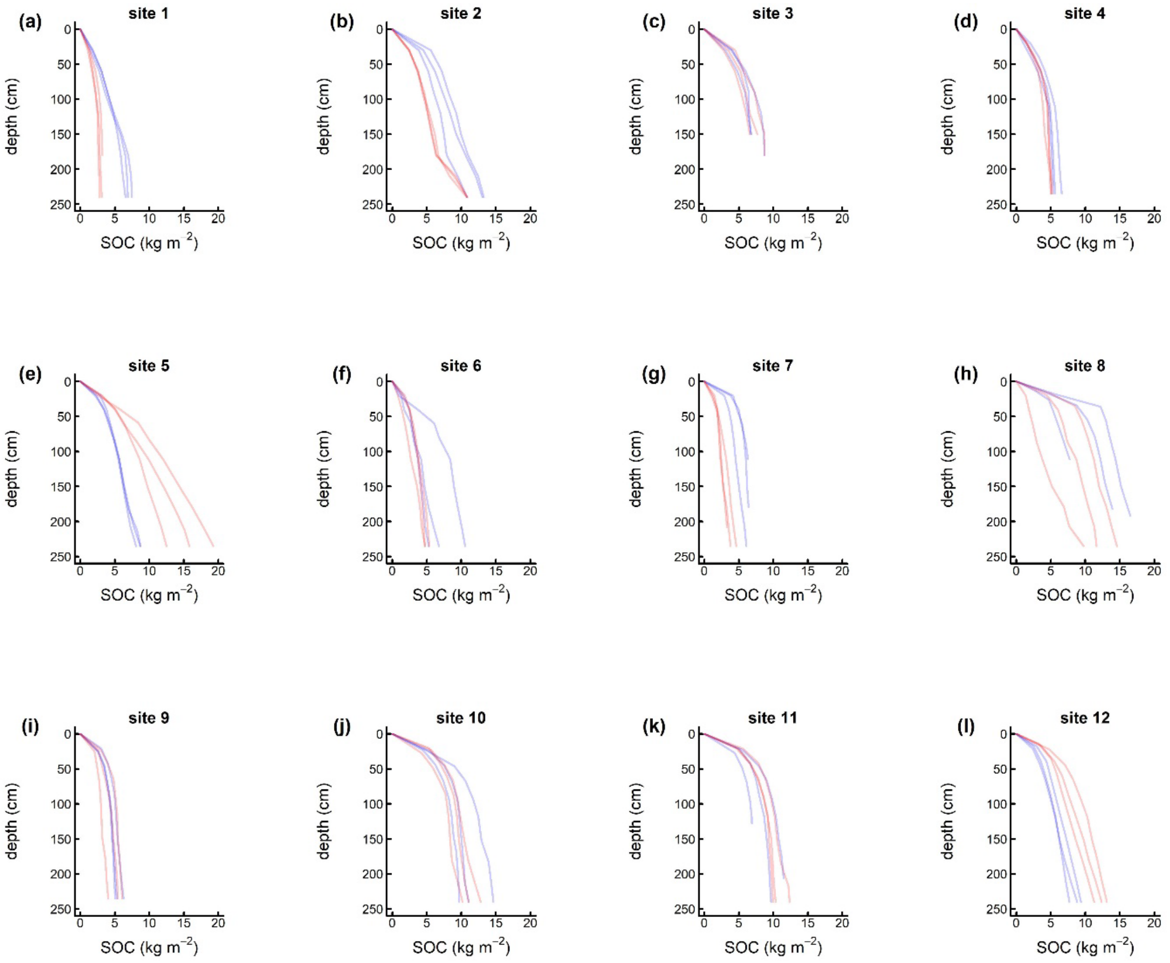


**Figure S1. Cumulative SOC stocks under switchgrass and shallow-rooted crops as a function of depth at 12 sites across the central and eastern USA**. Red lines show individual cores under shallow rooted crops and blue lines show cores under switchgrass.

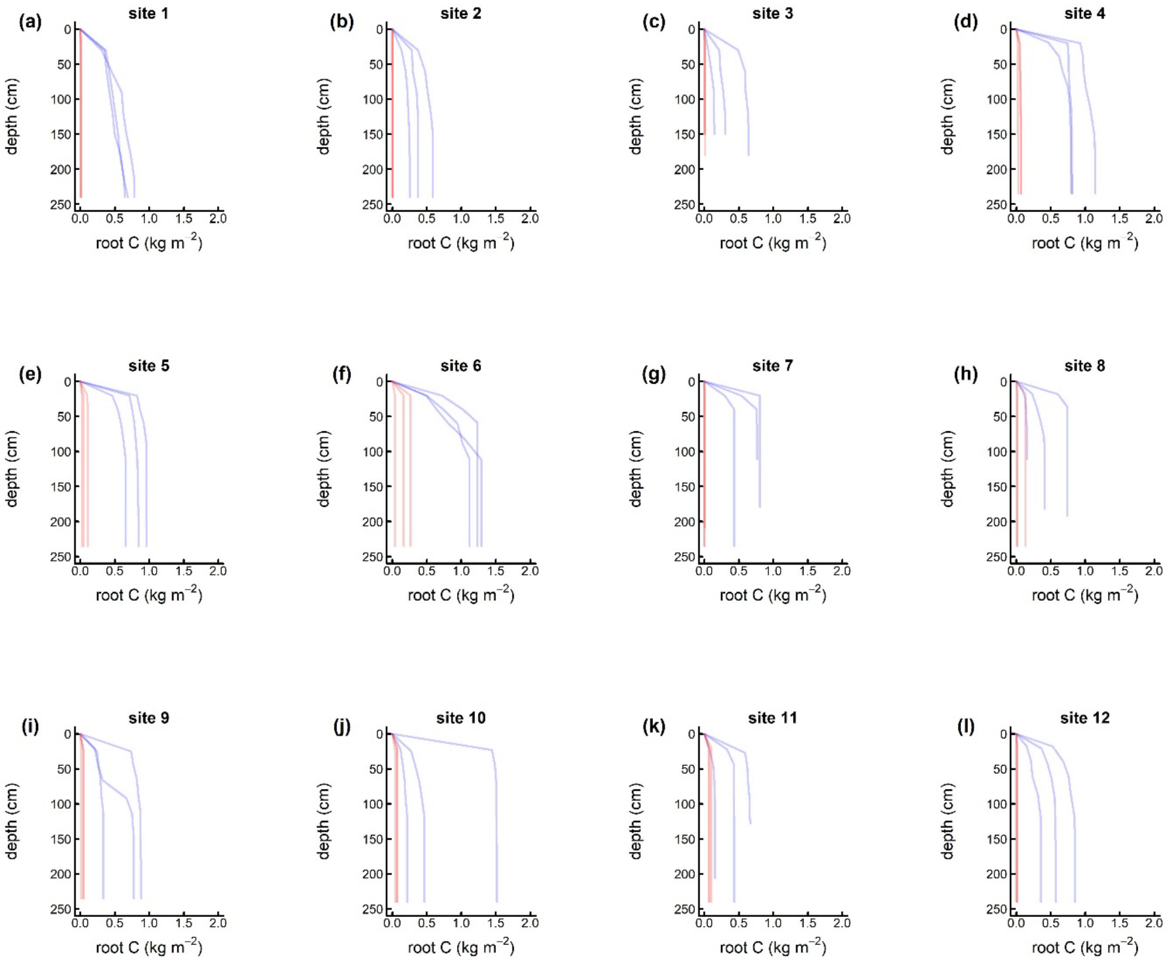


**Figure S1. Cumulative root C stocks under switchgrass and shallow-rooted crops as a function of depth at 12 sites across the central and eastern USA**. Red lines show individual cores under shallow rooted crops and blue lines show cores under switchgrass.

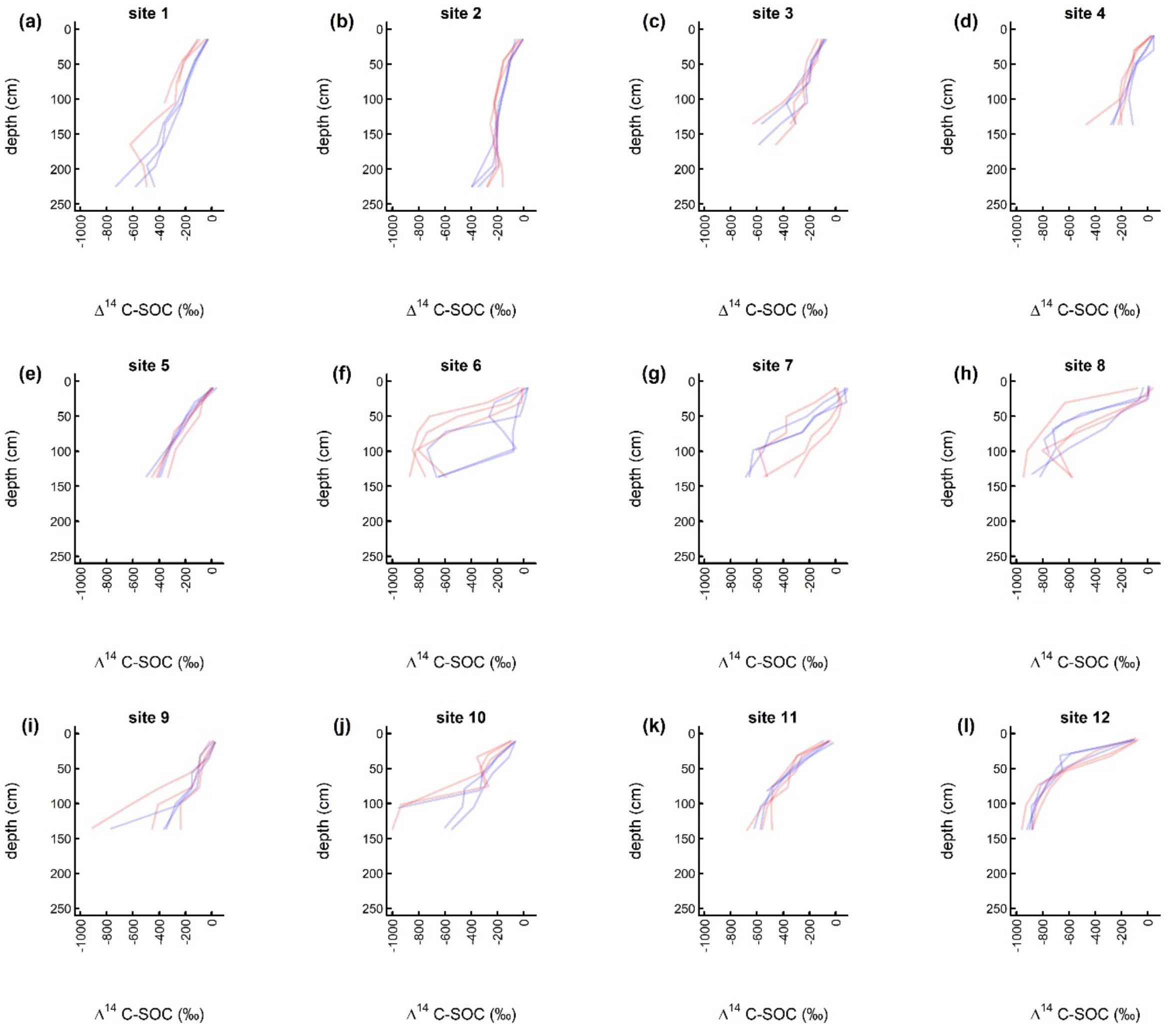


**Figure S3. Radiocarbon ($^{14}C$) content of SOC under switchgrass and shallow-rooted crops as a function of depth at 12 sites across the central and eastern USA**. Red lines show individual cores under shallow rooted crops and blue lines show cores under switchgrass.

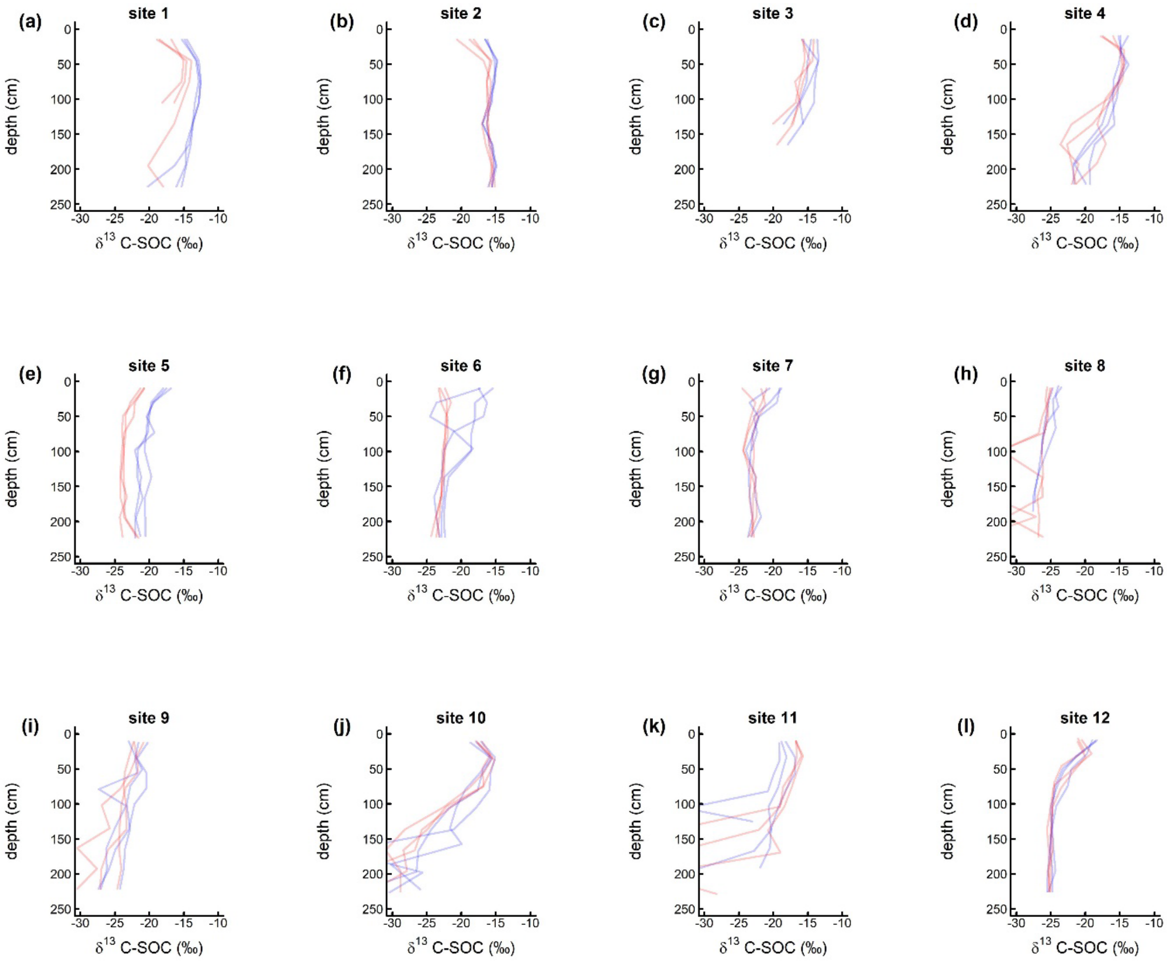


**Figure S1. $^{13}$C isotopic composition of SOC under switchgrass and shallow-rooted crops as a function of depth at 12 sites across the central and eastern USA**. Red lines show individual cores under shallow rooted crops and blue lines show cores under switchgrass.
